# Integration of ζ-deficient CARs into the *CD3-zeta* gene conveys potent cytotoxicity in T and NK cells

**DOI:** 10.1101/2023.11.10.565518

**Authors:** Jonas Kath, Clemens Franke, Vanessa Drosdek, Weijie Du, Viktor Glaser, Carla Fuster-Garcia, Maik Stein, Tatiana Zittel, Sarah Schulenberg, Caroline E. Porter, Lena Andersch, Annette Künkele, Joshua Alcaniz, Jens Hoffmann, Hinrich Abken, Mohamed Abou-el-Enein, Axel Pruß, Masataka Suzuki, Toni Cathomen, Renata Stripecke, Hans-Dieter Volk, Petra Reinke, Michael Schmueck-Henneresse, Dimitrios L. Wagner

## Abstract

I.

Chimeric antigen receptor (CAR)-reprogrammed immune cells hold significant therapeutic potential for oncology, autoimmune diseases, transplant medicine, and infections. All approved CAR-T therapies rely on personalized manufacturing using undirected viral gene transfer, which results in non-physiological regulation of CAR-signaling and limits their accessibility due to logistical challenges, high costs and biosafety requirements. Here, we propose a novel approach utilizing CRISPR-Cas gene editing to redirect T cells and natural killer (NK) cells with CARs. By transferring shorter, truncated CAR-transgenes lacking a main activation domain into the human *CD3*ζ *(CD247)* gene, functional CAR fusion-genes are generated that exploit the endogenous *CD3*ζ gene as the CAR’s activation domain. Repurposing this T/NK-cell lineage gene facilitated physiological regulation of CAR-expression and reprogramming of various immune cell types, including conventional T cells, TCRγ/δ T cells, regulatory T cells, and NK cells. In T cells, *CD3*ζ in-frame fusion eliminated TCR surface expression, reducing the risk of graft-versus-host disease in allogeneic off-the-shelf settings. *CD3*ζ-CD19-CAR-T cells exhibited comparable leukemia control to *T cell receptor alpha constant* (*TRAC*)-replaced and lentivirus-transduced CAR-T cells *in vivo*. Tuning of *CD3*ζ-CAR-expression levels significantly improved the *in vivo* efficacy. Compared to *TRAC*-edited CAR-T cells, integration of a Her2-CAR into *CD3*ζ conveyed similar *in vitro* tumor lysis but reduced susceptibility to activation-induced cell death and differentiation, presumably due to lower CAR-expression levels. Notably, *CD3*ζ gene editing enabled reprogramming of NK cells without impairing their canonical functions. Thus, *CD3*ζ gene editing is a promising platform for the development of allogeneic off-the-shelf cell therapies using redirected killer lymphocytes.

**Key points:**

- Integration of ζ-deficient CARs into *CD3*ζ gene allows generation of functional TCR-ablated CAR-T cells for allogeneic off-the-shelf use
- *CD3*ζ-editing platform allows CAR reprogramming of NK cells without affecting their canonical functions

## II. Introduction

The adoptive transfer of immune cells is a powerful tool to combat chronic diseases, such as cancer. Guiding lymphocytes to specifically bind and respond to antigens can be used to redirect the anti-tumor efficacy of cytotoxic T cells^1^ and natural killer (NK) cells^2^ as well as promote tissue-specific immunosuppression through regulatory T cells (Treg)^3,4^. To overcome the limitations associated with low frequencies of certain antigen-specific T cells in patients, gene transfer of chimeric antigen receptors (CAR) can be used to install the desired antigen-specificity to large numbers of cells needed for adoptive cell transfer and treatment success in severe disease. Autologous CAR-T cells are an approved treatment for B-cell malignancies, such as acute B-lymphoblastic leukemia^1,5^, B-cell lymphoma^6,7^ and multiple myeloma^8^.

The TCR/CD3-complex is the endogenous antigen-receptor in T cells. It consists of a TCRα and a corresponding TCRβ chain which engage antigenic peptides presented by MHC molecules, as well as the accessory proteins CD3γ, CD3δ, CD3ε and CD3ζ which transduce the TCR signal downstream. While all CD3 proteins are required for TCR/CD3 assembly, biosynthesis of CD3ζ is the rate-limiting step in TCR/CD3 complex formation^9^. Further, the intracellular domain of CD3ζ is sufficient to drive TCR-like activation in chimeric receptors^10,11^. Therefore, all clinically approved (second-generation) CARs use the intracellular domain of CD3ζ as their primary TCR-activation-like effector domain. CARs further comprise an extracellular antigen-binding domain, a hinge domain, a transmembrane domain and an additional intracellular co-stimulatory domain, such as CD28 or 4-1BB. CARs without a main activation domain do not induce cytotoxicity, but have been proposed to boost T cell function by providing co-stimulation^12^.

Most clinical CAR-T cell products are generated by transduction with viral vectors which randomly integrate their respective cargo into the genome and drive expression of the CAR through strong exogenous promoters, such as EF1α^5,8,13,16^. Positional effects and epigenetic silencing of exogenous expression cassettes have been linked to inconsistent CAR-expression levels^17,18^. While previous trials with virally transduced T cells have been safe overall^19^, gene transfer with (semi)-random integration poses the risk of insertional mutagenesis as highlighted by cases of clonal expansion after disruption of tumor suppressor genes *TET2*^20^ or *CBL*^21^ by integrated CAR provirus and the recent report of the development of CAR^+^ T cell lymphoma after treatment with products generated via PiggyBac transposase technology^22,23^.

Targeted gene transfer using gene editing can improve the consistency of redirected T cell products by predictable antigen receptor expression^17,24,25^. To this end, a programmable nuclease, such as CRISPR-Cas, is introduced into the T cells alongside a DNA repair template to exploit homology-directed DNA repair (HDR) for site-specific integration of the *CAR*-transgene. Multiple locations have been proposed to redirect T cells with CARs, including protein-coding genes such as *TCRα chain constant* (*TRAC*)^17,26–28^,*PDCD1 (*encoding PD-1)^27,29^ or *GAPDH*^30^ as well as intra-/extragenic genomic safe harbor (GSH) loci, such as the human AAV-integration site (*hAAVS1*)^29^ and *eGSH6*^18^, respectively. *TRAC* has emerged as the gold-standard for gene-edited CAR-T cells. One reason is the improved cell functionality associated with the temporary downregulation of the CAR after target engagement^17^. This mirrors the natural regulation of the human TCR and protects from overt differentiation and T cell exhaustion^17^. An additional advantage is that the integration of *CAR*-transgenes into *TRAC* disrupts the TCR/CD3-complex. This creates CAR^+^ TCR^-^ T cells which lack TCR-mediated allo-reactivity, thereby demonstrating a route towards safe application of CAR-T cells in allogeneic settings^31^.

In this study, we demonstrate virus-free CAR reprogramming via in-frame integration of truncated, CD3ζ-deficient *CAR*-transgenes (*trunc*CARs) into an early exon of the *CD3*ζ gene. Our knock-in strategy produces fusion genes composed of the exogenous *truncCAR*-transgene (encoding an antigen binder, a hinge, a transmembrane as well as a co-stimulatory domain but no main activation domain) and the endogenous *CD3*ζ gene. This reduces the required transgene size and exploits the endogenous *CD3*ζ promoter for physiological CAR regulation. *CD3*ζ gene editing can also be used for CAR reprogramming of regulatory T cells, TCRγ/δ T cells and most notably primary human NK cells which cannot be reprogrammed by *TRAC*-targeting.

## III. Material and methods

### Culture of primary cells

The study was performed in accordance with the declaration of Helsinki (Charité ethics committee approval EA4/091/19). Peripheral blood mononuclear cells (PBMC) were obtained from healthy donors via density gradient centrifugation from peripheral blood. T cells were enriched by magnetic cell separation (MACS) using CD3 microbeads and cultured in T cell medium, a 1:1 mixture of RPMI (Gibco) and Click’s (Irving) media supplemented with 10% fetal calf serum (FCS), IL-7 (10 ng/ml, CellGenix) and IL-15 (5 ng/ml, CellGenix). NK cells were enriched from the CD3-negative fraction using the NK isolation Kit (Miltenyi) and cultured in NK MACS Medium (Miltenyi) supplemented with 10% FCS, IL-2 (500 IU/ml) and IL-15 (5ng/ml).

### Genetic engineering

Targeted virus-free CAR integration was performed as recently described^32^. In short, human T or NK cells were transfected with precomplexed CRISPR-Cas9 ribonucleoproteins (RNP) and double-stranded DNA (dsDNA) to employ homology-directed DNA repair (HDR) (DNA/sgRNA Sequences: **Suppl. Table 1**). The dsDNA served as template for HDR and consisted of the (CAR/truncCAR) transgene flanked by 400 bp homology arms. Cells were resuspended in 20µl P3 Electroporation Buffer (Lonza) and electroporated with 1 µg HDR-template and 1.38 µl RNP consisting of synthetic modified single guide RNA (sgRNA, 100 μM, IDT), 15-50 kDa poly(L-glutamic acid)^33^ (100 μg/µl, Sigma) and recombinant SpCas9 protein (61 μM, IDT) in a 0.96:1:0.8 volume ratio using the 4D-Nucleofector (Lonza). Prior to electroporation, T cells were activated for 48 hours on αCD3/CD28-coated tissue culture plates and electroporated at a density of 5×10^4^ cells/μl buffer with the nucleofection program EH-115. Primary human NK cells were expanded in NK medium for 6-7 days and electroporated using program DA-100. The NK-92 line was electroporated at 2.5×10^4^ cells/μl with the program CA-137. Immediately after the electroporation, 100μl of the respective medium were added. 10min post-electroporation, T cells were transferred tnto medium supplemented with 0.5µM HDR-Enhancer v2 (IDT). For lentiviral (LV) controls, activated T cells were transduced 1 day post T cell isolation while being kept on αCD3/CD28 coated tissue culture well plates for another day. After editing, T cells were expanded in G-Rex 6-well plates (Wilson Wolf).

### Off-target analysis with CAST-Seq

The assay was performed using genomic DNA isolated from T cells 12 days after nucleofection as previously described^34,35^ (**Supplementary Methods**).

### Flow cytometry

Assessment of CAR^+^ rate, cytotoxicity, intracellular cytokine production, exhaustion, phenotype and CAR-regulation was performed on a Cytoflex LX device (Beckman Coulter) using the panels stated in **Suppl. Table 2** and as previously described^32^. Activation-induced cell death of Her2-CAR-T cells was assessed after stimulation with plate-bound anti-Fc antibody (10 μg/mL) (Jackson) by flow cytometry via staining for Annexin V Alexa Fluor® 647 stain (Biolegend) and 7AAD (Biolegend). NK cell degranulation was assessed after 4h of co-culture with target cells in the presence of Monensin A (1µM) and BV785-conjugated anti-CD107a antibody via flow cytometry. NK cell-mediated antibody-dependent cellular cytotoxicity (ADCC) was assessed after 16h of co-culture with CD20^+^ bGal^-^ Jeko-1 cells in the presence of anti-CD20 (Rituximab) or anti-bGal antibody (Invivogen).

### Live cell imaging

In vitro tumor control of HER2-CAR-T cells was assessed via live cell imaging of GFP-expressing cancer cells on an Incucyte device (Sartorius) (**Supplemental methods**).

### Animal experiments

The *in vivo* CAR-T cell potency studies were performed in accordance with the German animal welfare act and the EU-directive 2010/63. Animal studies 1 and 3 were approved by local authorities (Landesamt für Gesundheit und Soziales, LaGeSo Berlin, Germany) under the permission A0010/19. Study 2 was approved by the Lower Saxony Office for Consumer Protection and Food Safety – LAVES (permit number 16/2222). Detailed study protocols are included in the supplementary methods section. In brief, immunodeficient mice were infused with 0.5×10^6^ Nalm-6 cells (expressing *luciferase*) via tail vein injection. Four days later, 0.5×10^6^ or 1×10^6^ TCR-deficient CD19-CAR-T cells were infused intravenously. CAR-T cells were generated either via targeted integration of a CAR or a *trunc*CAR into the *TRAC* or *CD3*ζ gene, respectively, or by LV gene transfer and consecutive *TRAC*-knock-out (KO). Tumor burden was assessed using bioluminescence imaging. The staff carrying out the mice experiments were blinded for the T-cell conditions. Mice were sacrificed according to study protocol either at ethical endpoints (models 1+3) or five weeks after tumor inoculation (model 2) according to the respective animal study protocols.

### Data analysis, statistics and presentation

Flow cytometry data was analysed with FlowJo Software (BD). Prism 9 (GraphPad) was used to create graphs and perform statistics. Illustrations were created on BioRender.com.

### Data Sharing Statement

HER2-CARs were previously published^36^. Other CAR/HDR-templates and sgRNA sequences are provided in **Suppl. Table 1**. Plasmids encoding *CD3*ζ-HDR-templates will be distributed through Addgene.

## IV. Results

### Integration of truncated CD3ζ-deficient (trunc)CARs in CD3ζ enable reprogramming of T cells

We performed targeted delivery of a 1419bp-sized CD19-specific *trunc*CAR (CD19-IgG1-CD28) into *CD3*ζ (exon 2, beginning of intracellular domain) and *TRAC* (exon 1) using CRISPR-Cas (**Fig. 1a**). As additional control, we integrated a full-length 2015bp-sized CAR (CD19-IgG1-CD28-CD3ζ) into *TRAC* as recently described^32^. Transgene expression in primary human T cells was confirmed by flow cytometry (**Fig. 1b**). Like *TRAC-*editing, CAR integration into the *CD3*ζ gene ablated TCR/CD3 surface expression. In a VITAL-assay^37^, which monitors relative antigen-specific cytotoxicity, *TRAC*-edited *trunc*CAR-T cells did not elicit any antigen-specific cytotoxicity as expected due to the lack of a main activation domain (**Fig. 1c**). In contrast, *CD3*ζ-edited *trunc*CAR-T cells effectively lysed CD19^+^ cells similar to *TRAC*-edited T cells transfected with the full-length CAR (**Fig. 1c**), confirming the generation of functionally active *trunc*CAR-CD3ζ fusion protein after insertion of CAR moieties into the endogenous *CD3*ζ*-*gene.

**Fig. 1:**
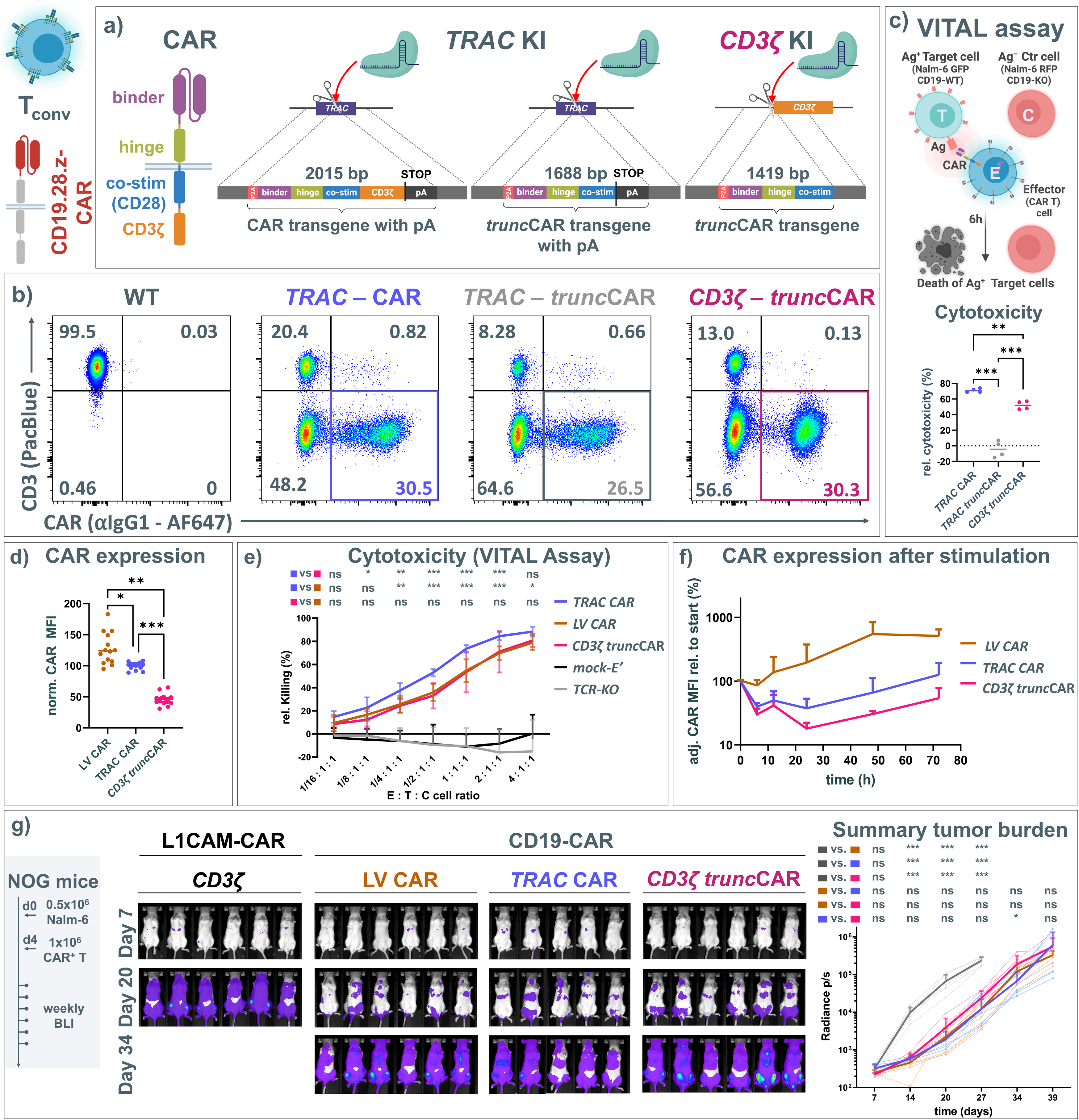
Integration of a truncated CD19-specific CAR into *CD3*ζ, but not *TRAC*, conveys cytotoxicity in conventional T cells toward CD19^+^ leukemia cells. (**a**) full-length second-generation CAR protein (left) and virus-free knock-in strategies to integrate a full-length CAR into *TRAC* or a truncated CAR (*trunc*CAR) into *TRAC* or *CD3*ζ. (**b**) Flow cytometry dot plots after knock-in. Transgene integration into *TRAC* or *CD3*ζ disrupts expression of the TCR/CD3 complex. (**c**) Relative cytotoxicity in co-culture with (CD19^+^) Nalm-6 target cells and CD19 knock-out Nalm-6 control cells (VITAL assay). Calculation of relative cytotoxicity according to formula stated in methods section. (n=2 biol. repl. each in 2 techn. repl.; ordinary one-way ANOVA followed by Holm-Šídák’s multiple comparison test with a single pooled variance). (**d-g**) Functional testing of *CD3*ζ *trunc*CAR, T cells in comparison to TRAC CAR and LV CAR-T cells. (**d**) Mean fluorescence intensity (MFI) determined by flow cytometry as a measure of cellular CAR-expression and normalized to each donor’s mean CAR MFI in the *TRAC* condition. (n = 7 biol. repl. each in 2-5 techn. repl.; mixed-effects analysis with Geisser-Greenhouse correction + Holm-Šídák’s multiple comparison test with individual variances computed for each comparison). (**e**) Relative cytotoxicity towards CD19^+^ cells assessed in a 6-hour VITAL assay. (mock-E’: mock-electroporated controls without RNP/HDR templates) (n=4 biol. repl. each in 1-3 techn. repl.; two-way ANOVA followed by Holm-Šídák’s multiple comparison test with a single pooled variance (**f**) Changes in CAR-expression levels (MFI normalized to start) after target cell encounter. (*TRAC* and LV in 4 biol. repl.; *CD3*ζ in 2 biol. repl.). (**g**) Acute lymphoblastic leukemia xenograft mouse model using luciferase-labeled Nalm-6 (CD19^+^) tumor cells. 4 days post Nalm-6 administration, 1×10^6^ cryopreserved, 14-day expanded TCR-deleted CAR^+^ T cells were injected systemically. Tumor burden was assessed via bioluminescence imaging (BLI). (n=5-6; 2-way ANOVA with Geisser-Greenhouse correction of log-transformed BLI data followed by Holm-Šídák’s multiple comparison test, with individual variances computed for each comparison). Asterisks in this and all further figures represent different p-values calculated in the respective statistical tests (ns: p > 0.05; *: p < 0.05; **: p < 0.01; ***: p < 0.001).

### CD3ζ-truncCAR and TRAC-CAR-T cells have comparable CAR-regulation and anti-leukemia activity

We next compared CD19-CAR-expression levels and anti-leukemia potential of *CD3*ζ-*trunc*CAR-T cells, *TRAC*-CAR-T cells and lentivirus-transduced (LV) *TRAC*-KO CAR-T cells *in vitro*. CAR-expression levels in *CD3*ζ-*trunc*CAR-T cells were lower than in *TRAC*-integrated and LV counterparts (**Fig. 1d**). Compared to *TRAC*-CAR-T cells, *CD3*ζ-*trunc*CAR-T cells and LV CAR-T cells displayed slightly reduced dose-dependent killing in a 6-hour VITAL assay (**Fig. 1e).** Upon CD19^+^ Nalm-6 target cell engagement, *CD3*ζ*-trunc*CAR and *TRAC*-CAR-T cells downregulated the CAR for 12-24 hours before returning to their relative baseline levels (**Fig. 1f**). In contrast, LV CAR-T cells upregulated CAR-expression in response to stimulation and exceeded their baseline levels after 48 hours. Previous studies demonstrated that physiological control of CAR-expression in the *TRAC* locus enhances their anti-tumor performance *in vivo*^17^. Therefore, we evaluated the anti-tumor efficacy of the differently engineered T cells (LV, *TRAC*, *CD3*ζ*-trunc*CAR) in two independent, blinded xenograft models of acute lymphoblastic leukemia using immunodeficient mice. In both experiments, 0.5×10^6^ luciferase-labeled CD19^+^ Nalm-6 tumor cells were administered systemically prior to the infusion of TCR-deficient CAR-T cells four days later. In mouse model 1 (**Fig. 1g**), mice received 14-day expanded cryopreserved CAR-T cells at a dose of 1×10^6^ CAR^+^ cells. All three CAR-T treatments slowed tumor growth to a similar extent (control: L1CAM-CAR^38^). *In vivo* efficacy was also observed in mouse model 2 (**Suppl. Fig. 1**). Here, fresh, 18-day expanded CAR-T cells were administered at a dose of 0.5×10^6^ CAR^+^ cells.

### Increasing CAR-expression from CD3ζ improves IL-2 production and anti-tumor efficacy

We hypothesized that the lower short-term cytotoxicity of CD3ζ-*trunc*CAR-T cells is caused by the lower amounts of CAR molecules available for synapse formation. Optimization of the 2A-cleavage peptide by the addition of a GSG-linker has been shown to increase protein expression in multi-cistronic transgenes^39,40^. In the *CD3*ζ*-trunc*CAR condition, an optimized GSG-P2A (**Fig. 2a**) increased CAR-expression even above the *TRAC*-CAR condition (**Fig. 2b**). Indeed, this modification increased CAR-mediated cytotoxicity (**Fig. 2c**) and intracellular cytokine production to levels similar to *TRAC*-CAR-T cells (**Fig. 2d, Suppl. Fig. 2**). We next evaluated the impact of the different CAR-expression levels during repeated leukemia challenges (**Fig. 2e-h**) which were performed once per week at a CAR^+^ T cell to tumor cell ratio of 1:1. After serial co-culture, all three conditions retained their physiological CAR expression dynamics, but basal CAR-expression did not differ anymore between *CD3*ζ*-trunc*CAR^GSG^ and *TRAC*, while the original *CD3*ζ*-trunc*CAR cells still showed lower CAR-expression (**Fig. 2e**). Interestingly, all three conditions showed similar cytotoxicity (**Fig. 2f**) and proliferation (**Fig. 2g**). *CD3*ζ-edited conditions displayed slightly lower expression of inhibitory markers in the CD8 compartment after serial leukemia re-challenges (**Fig. 2h; detailed analysis in Suppl. Fig. 3**). Serial co-culture resulted in a similar shift towards a more differentiated phenotype in all three conditions (**Suppl. Fig. 4a**) with a trend towards a CD8 polarization in the *CD3*ζ-*trunc*CAR^GSG^ condition (**Suppl. Fig. 4b**). Of note, the differences in cytokine production were preserved (**Suppl. Fig. 4c**). Finally, we assessed the *in vivo* anti-tumor efficacy of the three conditions in a Nalm-6 mouse model (**Fig. 2i**). *TRAC*-CAR-T cells and *CD3*ζ*-trunc*CAR both result in a similarly prolonged, statistically significant survival compared to mock-electroporated T cells. Expression-tuned *CD3*ζ*-trunc*CAR^GSG^-T cells showed the highest survival benefit which was statistically significant to the other treatment groups. *Ex vivo* expansion of CAR-T cells was reduced to 6 days due to a preferable phenotype with a high proportion of central memory (T_CM_) and naïve-like (T_N_) cells as well as a physiological CD4/CD8 ratio at this time point (**Suppl. Fig. 5**).

**Fig. 2:**
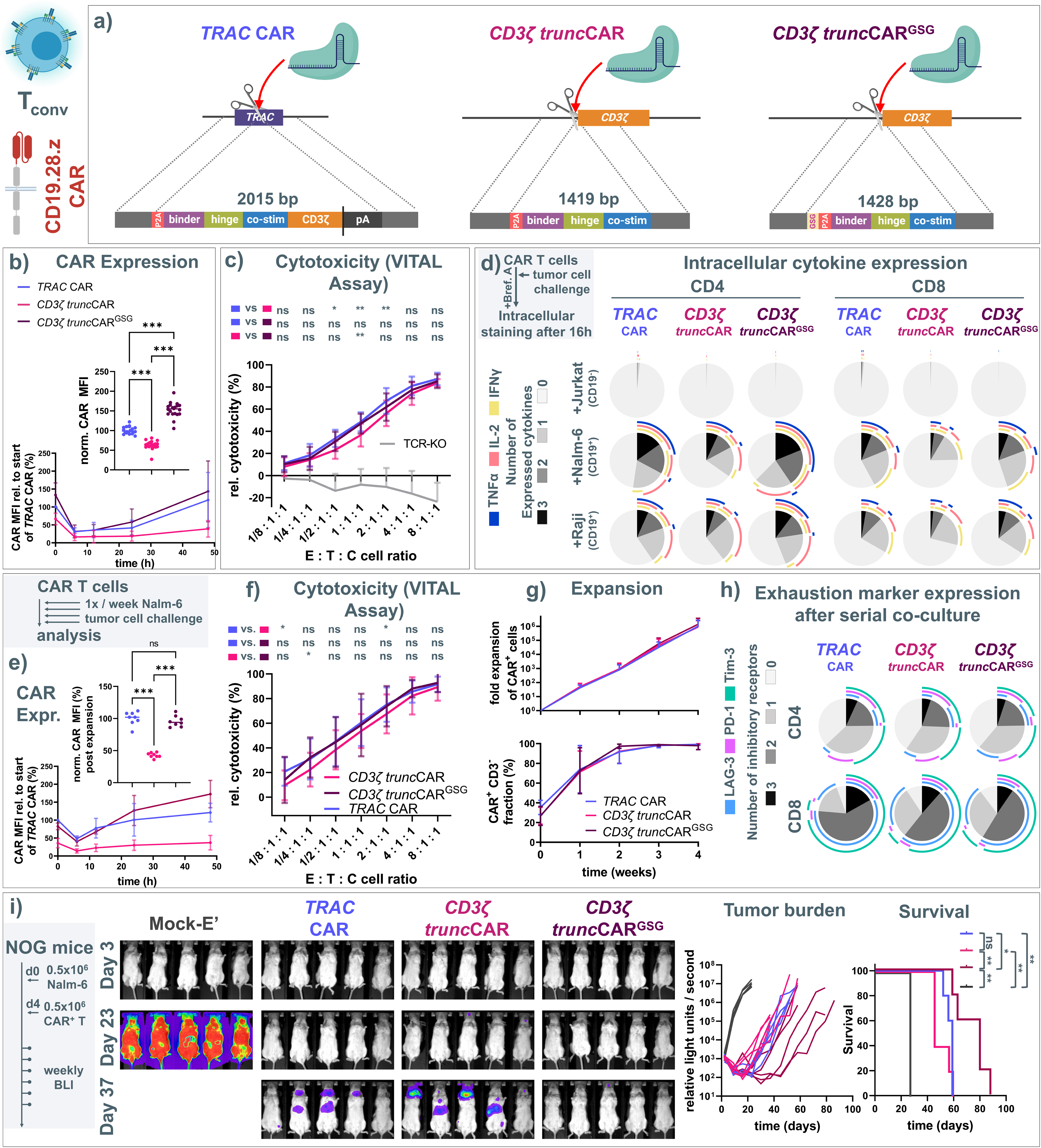
Evaluation of an optimized *CD3*ζ *trunc*CAR transgene and its impact on CAR-T cell function *in vitro*. (**a**) dsDNA templates for targeted delivery of a CAR or *trunc*CAR respectively into *TRAC* (left) or *CD3*ζ (middle), as in Fig. 1a, and for targeted delivery of a GSG-P2A-linker-modified *trunc*CAR into *CD3*ζ (right). (**b**) Top: Mean fluorescence intensity (MFI) determined by flow cytometry at steady (n=4 biol. repl. in 4-6 techn. repl. in two independent experiments, data normalized to mean of *TRAC* for each donor; mixed-effects analysis with Geisser-Greenhouse correction followed by Holm-Šídák’s multiple comparison test, with individual variances computed for each comparison). Bottom: dynamics of CAR MFI after CAR-stimulation using CD19^+^ Nalm-6 tumor cells. (n = 3-4 biol. Replicates in 1-2 techn. replicates). (**c**) Relative cytotoxicity assessed in a 6-hour VITAL assay (similar to Fig. 1c, n=4 biol. repl. in 3 techn. repl.; two-way ANOVA followed by Holm-Šídák’s multiple comparison test with a single pooled variance.). (**d**) Cytokine expression in CAR^+^ cells in response to control (CD19^-^) cell or target (CD19^+^) cell encounter (n=3 biol. repl.). (**e-h**) CAR-T cell re-challenge in serial co-cultures with Nalm-6 target cells. (**e**) Top: CAR MFI normalized to *TRAC* condition at steady state (n=2 biol. repl. in 4 techn. repl.; statistics as in b). Bottom: dynamics of CAR MFI after target cell engagement (n = 2-4 biol. repl. in 1-2 techn. repl.). (**f**) 6-hour VITAL assay. (n=3 biol. repl. in 3-4 techn. repl.; two-way ANOVA followed by Holm-Šídák’s multiple comparison test with a single pooled variance.). (**g**) Top: relative expansion of CAR^+^ T cells (top); Bottom: CAR^+^ frequency within T cell products. (n= 4 biol. repl.). (**h**) Cell surface expression of inhibitory receptors (LAG-3, PD-1, TIM-3; means of n=4 bio. repl.). (**i**) *In vivo* CAR-T cell efficacy tested in Nalm-6 acute lymphoblastic leukemia xenograft mouse model (n=5-6 mice/group; multiple log-rank tests).

### Tightly controlled HER2-CAR-expression from CD3ζ avoids antigen-independent differentiation and protects from activation-induced cell death

CAR-T cell therapies have also been developed for solid tumor-associated antigens, such as HER2^41–43^. To test our *CD3*ζ-editing platform in this setting, we generated HER2-specific CAR-T cells via integration of a *trunc*CAR into *CD3*ζ. As controls, we integrated of a full-length CAR into *TRAC*, or into the safe-harbor locus *hAAVS* driven by an exogenous LTR/EF1α-promoter. *CD3*ζ-edited HER2-*trunc*CAR-T cells demonstrated the lowest CAR-expression level (**Suppl. Fig. 6a**). *TRAC*-edited T cells displayed higher CAR-expression than the LTR/EF1a-driven CAR from the *hAAVS1* locus. Phenotype analysis demonstrated antigen-independent differentiation in an expression level dependent manner (**Suppl. Fig. 6b**). *TRAC*-HER2-CAR-T cells expressed the highest levels of inhibitory receptors PD-1, Lag-3 and Tim-3 after two weeks expansion (**Suppl. Fig. 6c**). In contrast, *CD3*ζ-HER2-*trunc*CAR-T cells displayed differentiation status and exhaustion marker profiles mirroring the CAR**^-^** T cell fraction which indicates reduced or absent tonic signaling. Further, *CD3*ζ-edited HER2-*trunc*CAR-T cells demonstrated lower expression of markers for early apoptosis than *TRAC*- or *AAVS1*-edited CAR-T cells after CAR stimulation using plate-bound antibody, highlighting their reduced propensity for activation-induced cell death (**Suppl. Fig. 6d**). Finally, *CD3*ζ-*trunc*CAR-T cells showed identical cytotoxicity toward three different HER2**^+^** tumor cell lines when compared to *TRAC*-HER2-CAR-T cells (**Suppl. Fig. 6e**). Therefore, *CD3*ζ gene editing may also serve as a platform to redirect T cells towards solid cancers.

Importantly, off-target assessment with CAST-Seq^34^ indicated high precision of the CRISPR-Cas9-mediated *CD3*ζ*-*targeting. The analysis did not reveal any chromosomal translocations, only the expected on-target aberrations including a very rare 15 Mb deletion between CD3ζ and a potential off-target site located on the same chromosome (**Suppl. Fig. 7**).

### CD3ζ-targeting allows redirection of more immune cell types than TRAC-editing

Non-conventional T cells and natural killer (NK) cells have emerged as important CAR carriers for adoptive cell transfer^2,3,44–46^. To test the suitability of *CD3*ζ-editing for different cell therapy applications, we compared *CD3*ζ*-trunc*CAR and *TRAC*-CAR integration in TCR_γ/δ_ T cells, regulatory T (T_reg_) cells and primary NK cells (**Fig. 3**). Like *TRAC*, *CD3*ζ is expressed in all TCR_α/β_ T cells and gene editing of the respective loci led to similar frequencies of HLA-A2-specific CARs in Treg cells (**Fig. 3a, Suppl. Fig. 8a**). Furthermore, *CD3*ζ is expressed in other immune cells which do not express *TRAC* and should therefore not be targetable by in-frame *TRAC* integration, notably TCR_γ/δ_ T cells and natural killer (NK) cells. Of note, *TRAC*-editing in TCR_γ/δ_ T cells resulted in substantial CAR^+^ fractions, suggesting mRNA transcription of the *TRAC* gene in TCR_γ/δ_ T cells at a steady state (**Fig. 3b, Suppl. Fig. 8b**). As expected for NK cells, *trunc*CAR integration into *CD3*ζ, but not *TRAC*, led to detectable CAR-expression. Therefore, *CD3*ζ gene editing may serve as a universal approach to redirect different conventional and non-conventional T cells as well as NK cells with CARs (**Fig. 3c**).

**Fig. 3:**
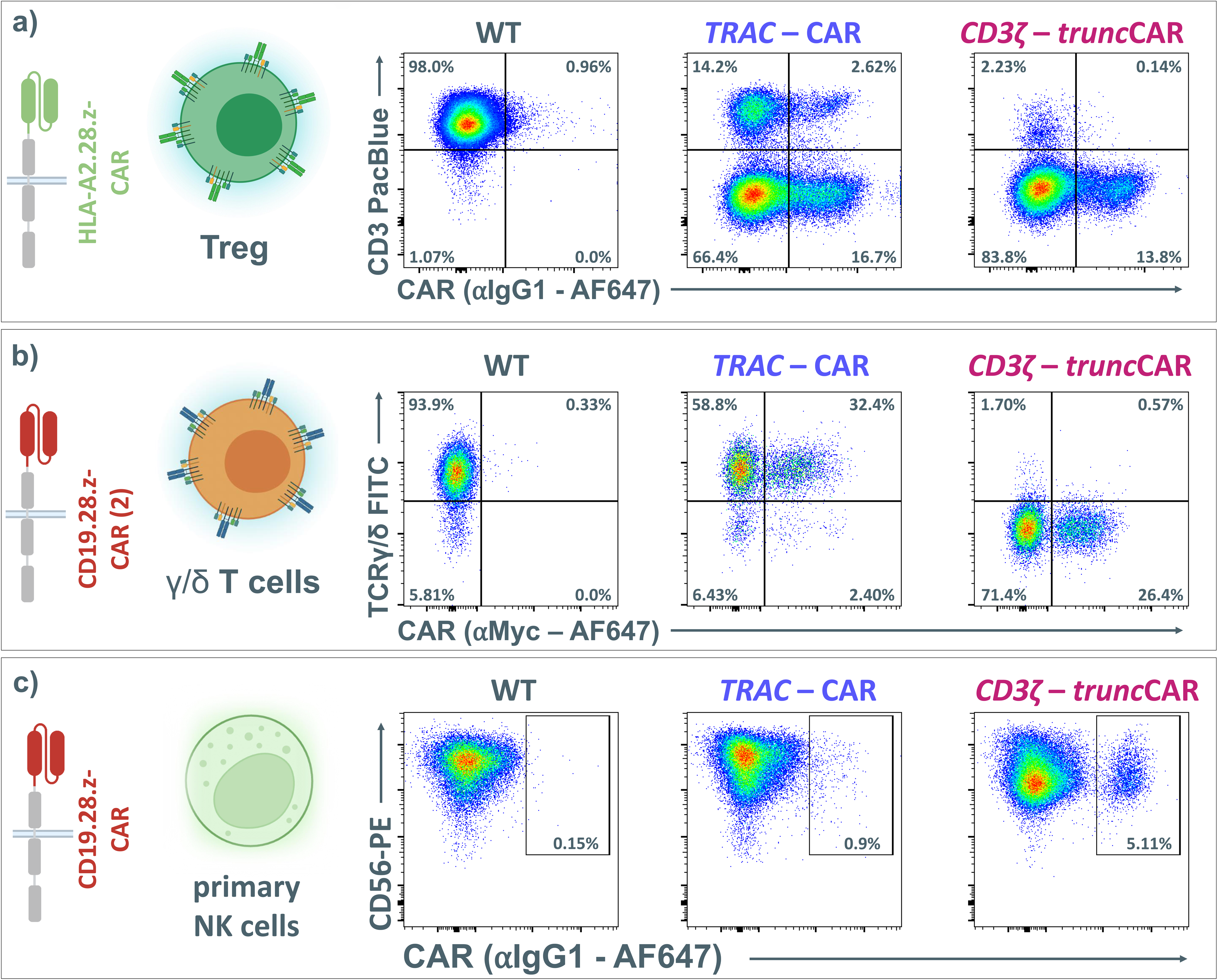
*CD3*ζ *trunc*CAR integration facilitates CAR-expression in different non-conventional T cell subtypes and NK cells. (**a**) HLA-A2 CAR integration in regulatory T cells. (**b**) CD19-CAR integration in TCR_γ/δ_ T cells. *TRAC* integration generates CAR^+^/ TCR_γ/_ ^+^ double positive T cells. (**c**) Integration of a CD19-CAR in primary human NK cells.

### CD3ζ-KO does not impede canonical functions of primary NK cells

In NK cells, CD3ζ is an adapter protein which assembles with activating killer-cell immunoglobulin-like receptors (KIR) and Fc-receptors, such as CD16^46^. NK cells continuously integrate inhibitory and activating signals shifting toward target cell killing when sensing enhanced KIR-activation (e.g. by increases in stress- and cancer-associated markers like Mic-a/b) or if CD16 triggers ADCC.Our knock-in approach impedes the expression of free CD3ζ-protein, which could potentially impair NK cell activation and disturb canonical NK functions. To investigate these potential downsides, we disrupted *CD3*ζ in primary human NK cells, either via CRISPR-Cas9-mediated KO or via *CD3*ζ-GFP-reporter knock-in that disrupts *CD3*ζ (**Suppl. Fig. 9a**). Measuring cytotoxicity (**Suppl. Fig. 9b**) and degranulation (**Suppl. Fig. 9c)** in simple co-cultures, we did not observe major differences regarding missing-self activation, cancer-directed activation, and allo-reactivity. Importantly, gene editing of *CD3*ζ did not alter CD16 expression. (**Suppl. Fig. 10a**). We also did not detect differences in anti-CD20-antibody-induced CD16-mediated ADCC towards the CD20^+^ cell line Jeko-1 (**Suppl. Fig. 9d**) which is partially resistant to NK cell cytotoxicity (**Suppl. Fig. 10b**).

### CD3ζ-truncCAR knock-in conveys cytotoxicity in primary NK cells and NK-92 cells

Using PBMC-derived NK cells, we next sought to characterize and compare *CD3*ζ-truncCAR-NK cells with LV-transduced NK cells (**Fig. 4**). *CD3*ζ-truncCAR knock-in rates remained below 10% and were thus considerably lower than in T cells (**Fig. 4a**). However, using the same LV as for the T cells, CAR transduction rates were in the same range despite a high multiplicity of infection (MOI=5). CAR MFI did not significantly differ between the conditions (**Fig. 4b**). Both conditions, but not a *TRAC*-CAR knock-in control, showed dose-dependent CAR-mediated killing with a trend towards superiority of the *CD3*ζ-truncCAR-NK cells in a VITAL assay, an internally controlled co-culture assay which is less biased by the NK cells’ CAR-independent (background-) killing (**Fig. 4c**). Analysis of the degranulation marker CD107a further confirmed CAR-mediated activation of CAR^+^ NK cells when co-cultured with CD19-expressing allogeneic B cells. However, this effect was only statistically significant for the *CD3*ζ-truncCAR condition (**Fig. 4d**). As for *CD3*ζ-KO cells (**Suppl. Fig. 9**), ADCC towards the CD20^+^ cell line Jeko-1 was not altered for *TRAC*, LV or *CD3*ζ-*trunc*CAR-NK cells compared to mock-electroporated (wildtype) NK cells (**Fig. 4e**). Thus, *CD3*ζ gene editing may be used to redirect primary NK cells with CARs while retaining their canonical functions. The NK-cell-derived cancer cell line NK-92 has been used as the cell source for CAR-NK therapy in clinical trials^47^. The use of immortal cell lines does not require high CAR integration rates because the edited cells can be enriched prior to a potentially unlimited expansion. To test the feasibility of our approach in NK-92 cell, we generated CD19-specific *CD3*ζ-*trunc*CAR-NK cells and *hAAVS1*-CAR-NK-92 cells as controls (**Fig. 4f**). CAR^+^ NK-92 cells were enriched via MACS. Compared to *hAAVS1*, *CD3*ζ*-trunc*CAR-NK-92 cells displayed higher CAR-mediated cytotoxicity (**Fig. 4g**) and superior (CAR-independent) missing-self activation towards the MHC-I-deficient cell line K562 (**Fig. 4h**). As NK-92 cells do not express CD16, ADCC was not studied.

**Fig. 4:**
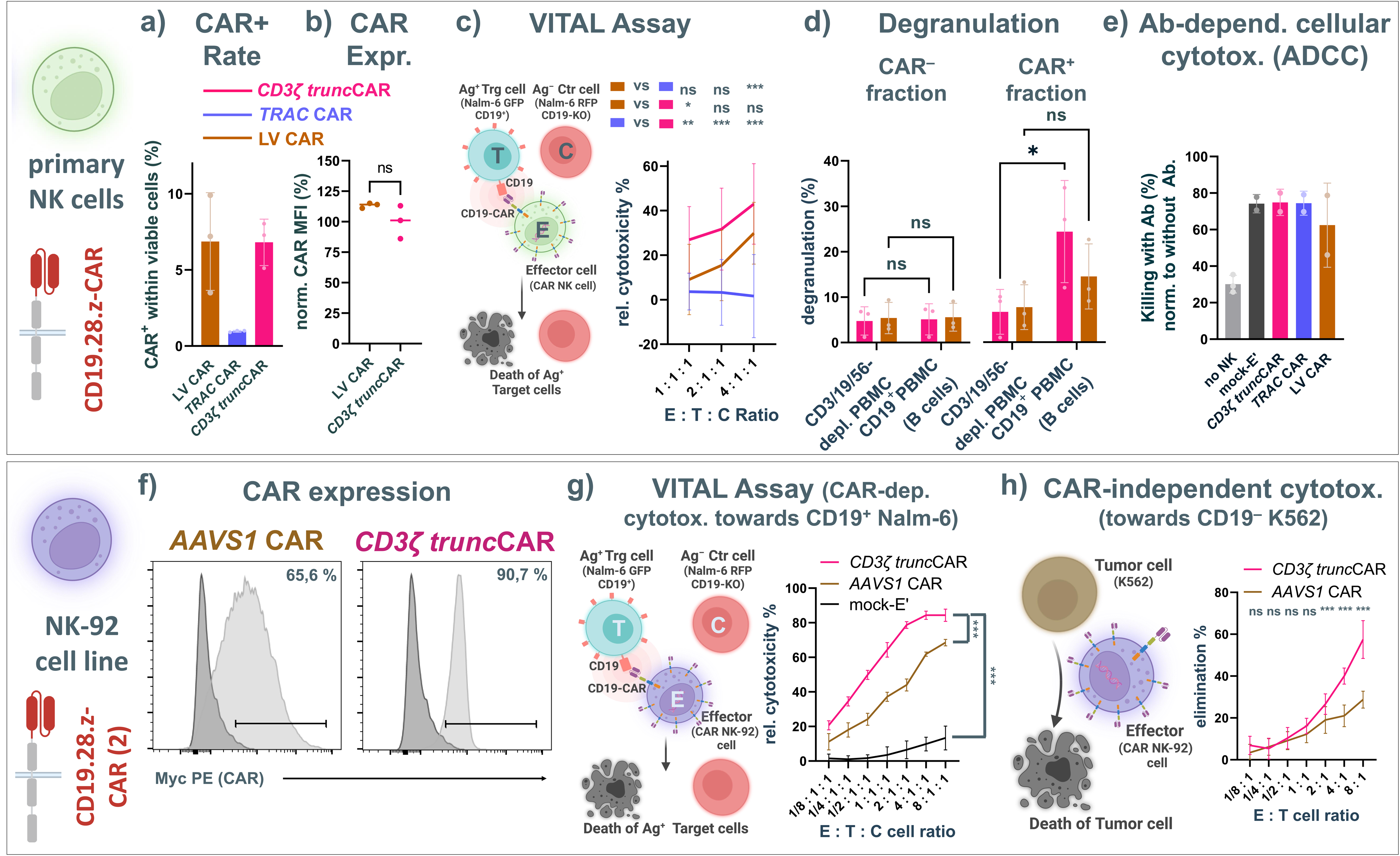
*CD3*ζ-editing enables redirection of NK cells with CARs and does not impede canonical NK cell functions *in vitro*. CAR editing in primary NK cells via LV CAR transfer, *TRAC*-CAR or *CD3*ζ*-trunc*CAR integration: (**a**) CAR^+^ frequencies after editing (n=3 biol. replicates); (**b**) mean CAR-expression in CAR^+^ cells (n=3 biol. replicates; Student’s t test.); (**c**) CAR-dependent cytotoxicity detected in a VITAL assay (data normalized to mock-electroporated (wildtype) NK cells; n=3 biol. repl. each in 3-4 techn. repl.; 2-way ANOVA followed by Tukey’s multiple comparison test with a single pooled variance); (**d**) Degranulation as indicator of NK effector function via flow cytometric detection of CD107a (n=3 biol. repl.; two-way ANOVA followed by Holm-Sidak’s multiple comparison test with a single pooled variance); (**e**) antibody-dependent cellular cytotoxicity (ADCC) of primary (CAR) NK cells against CD20^+^ bGal^-^ Jeko-1 cells. Bars represent killing for each condition in the presence of the CD20-targeting monoclonal antibody Rituximab (0.5µg/ml) normalized to the respective condition without supplemented Rituximab (n=2 biol. repl.); (**f-h**) CD19-CAR (2) transfer to NK-92 cells via *AAVS1* integration of a CMV promotor-controlled, full-length CAR or *CD3*ζ integration of a *trunc*CAR. CAR^+^ fractions were enriched using MACS. (**f**) CAR-expression in flow cytometry histograms. (**g**) CAR-dependent cytotoxicity in a 4-hour VITAL-assay (n=6 techn. repl.; two-way ANOVA with Tukey’s multiple comparison test with a single pooled variance. (**h**) CAR-independent cytotoxicity towards the MHC I deficient, CD19^-^ K562 (control) cell line (n=15 techn. repl.; two-way ANOVA followed by Holm-Šídák’s multiple comparison test with a single pooled variance).

## V. Discussion

Here, we propose a novel strategy for site-specific *CAR* gene transfer to T and NK cells. Truncated *CAR*-transgenes lacking a TCR-like effector domain were precisely inserted into the *CD3*ζ gene. Via in-frame integration, a complete CAR fusion gene (comprising an exogenous truncated *CAR*-transgene and the endogenous *CD3*ζ gene) is formed resulting in surface expression of functional CAR proteins. In T cells, this prevents TCR/CD3 complex assembly and brings the CAR under the transcriptional regulation of the *CD3*ζ gene. Despite its function as a signal transducer of activating NK cell receptors, *CD3*ζ can be edited to generate functional CAR-NK cells without affecting their canonical functions.

First clinical trials demonstrated that TCR-deleted allogeneic CAR-T cells can induce remissions in heavily pre-treated B-ALL and B-lymphoma patients, but additional gene editing was needed to circumvent immunological barriers of HLA-mismatches between CAR-T cell donor and patient^50–52^. Therefore, *CD3*ζ-editing would benefit from other modifications to improve the efficacy of allogeneic CAR-T cells^52,53^. Future studies may investigate the combination of *CD3*ζ-editing with additional KOs to improve functionality^54,55^, safety^56^ as well as persistence^57,50,58^ of allogeneic T and NK cells. Although *CD3*ζ-editing can be used for both T and NK cells, the respective edits required to improve the functionality of NK cells^59,60^ may differ to the ones proposed for T cells^55,61^. Finally, complex editing may require the combination of nuclease-assisted gene transfer with other gene silencing modalities such as base editing^62,63^ to reduce the risk for genomic rearrangements with unknown biological impact^52,61,64^.

This study is the first to demonstrate non-viral CRISPR-Cas-mediated knock-in for functional reprogramming of primary human NK cells with CARs. In comparison to CAR-T cells, CAR-NK cells have a favorable safety profile as they lack alloreactivity, do not persist long-term and show a reduced incidence of severe cytokine release syndrome and neurotoxicity^2^. CAR-NK cells can be combined with monoclonal antibodies for synergistic activity when targeting heterogenous tumors. For example, the CD19-specific CAR-NK cells generated by *CD3*ζ-editing (**Fig. 3-4**) may be combined with rituximab to overcome antigen-escape and relapse by CD19-negative cancer cells. Prior to testing in suitable *in vivo* models and future clinical translation, the efficacy of non-viral reprogramming of primary NK cells should be further increased, for example by using pharmacological enhancers^32^ and/or end-modified ssDNA donor templates^65^.

The *CD3*ζ locus is a novel CAR-integration site which shares features and advantages with the *TRAC* knock-in^17,26^. Like *TRAC*-, *CD3*ζ*-*editing causes TCR-ablation. Furthermore, the CAR’s CD3ζ-domain cannot rescue TCR/CD3-expression in *CD3*ζ-KO T cells^66^. Together, this avoids the risk of alloreactivity in TCRα/β^+^ T cells. Despite efficient *CD3*ζ-editing, residual TCR^+^ T cells must be depleted prior to allogeneic application to further minimize the risk for GvHD^52,67^. Further, the physiological TCR-like CAR-downregulation after antigen-engagement (achieved via *TRAC*- or *CD3*ζ-integration) may enable transient resting, preventing terminal differentiation and exhaustion^17,68^. When considering autologous manufacturing, transgene expression from TCR/NK-cell lineage genes, such as *TRAC* or *CD3*ζ, provides a safety advantage because it should prevent the inadvertent CAR-expression in B-cell leukemic blasts which can lead to CD19-antigen masking and B-ALL relapse^69^.

Deleterious mutations of *CD3*ζ have been found to be a cause for severe combined immunodeficiency, and NK cells obtained from these patients were found to be hypo-responsive in tumor co-cultures and after CD16 stimulation^70,71^. However, in this study, *CD3*ζ disruption did not result in any changes in ADCC, cytotoxicity or degranulation in primary human NK cells (**Fig. 4**). This is in line with previous findings that the signal transducer FcRγ compensates CD3ζ-loss after KO to enable ADCC^72^.

The serendipitous finding that *TRAC* knock-in led to CAR-expression in TCRγ/δ+ T cells (**Fig. 3**) warrants future investigations. The gene locus for TCRα and TCRδ is interconnected with *TRAC* being located downstream of the TCRδ constant (*TRDC*) gene^73^. Our results indicate that some TCRγ/δ^+^ T cells express the TCRα-chain at least from one allele.

CAR-expression level influences CAR-T cell performance, differentiation and exhaustion in pre-clinical and clinical settings^17,74,75^. For viral gene transfer, CAR density may be modulated by variation of viral titers, aiming for different transgene copy numbers, as well as different promoters^76^ or transgene designs^74^. Exogenous promoters required for CAR-expression after random integration can cause unphysiological CAR up-regulation after antigen-encounter (**Fig. 2c**) which can lead to cellular exhaustion^17^. The rules of transgene expression from genome-encoded genes, such as *CD3*ζ, via their endogenous promoters is poorly understood thus far. Transgene expression can be affected by the promoter as well as the 5’- or 3’-UTR and this could contribute to differences detected between *CD3*ζ and *TRAC*. However, we have also observed transgene-dependent differences (CD19-CAR vs HER2-CAR, see **Suppl. Fig. 7**), that were locus-dependent and warrant further investigation. We show that basal CD19-CAR-expression can be increased by insertion of a GSG-linker before the 2A-self-cleavage peptide (**Fig. 2**). Modulation of both, steady-state and dynamic CAR regulation, may impact the activation threshold of the CAR-T cells. Increasing the CAR activation threshold may reduce on-target off-tumor toxicity when targeting tumor-associated antigens upregulated in the tumor but not completely absent in normal tissue^77^. While low CAR-expression in *CD3*ζ-*trunc*CAR-T cells was sufficient to trigger cytotoxicity, it was associated with lower cytokine production after antigen-engagement and lower anti-leukemia activity *in vivo* (**Fig. 2**). Of note, all CARs used in this study employed the CD28 co-stimulatory domain. Future studies should revisit the contribution of other co-stimulatory domains in CAR-T/NK cells created by *CD3*ζ-gene editing to select the most efficacious CAR-version for the targeted disease.

## VI. Acknowledgements

We would like to express our gratitude to the following individuals for their valuable contributions: Laila Hassan (Charité, Berlin, Germany; † deceased) for her technical assistance with the experiment presented in Fig. 1. Silke Schwiebert from Annette Künkele Lab (Charité, Berlin, Germany) for her assistance with lentivirus preparation. Alina Pruene and Tobias Braun (Medizinische Hochschule Hannover, Hannover, Germany) for their support in animal model 2. Amanda Roswell-Shaw and Daniel Wang (Baylor College of Medicine, Houston, USA) for their assistance with HER2-CAR-T cell co-cultures. Geoffroy Andrieux (from the Institute of Medical Bioinformatics and Systems Medicine, Medical Center-University of Freiburg) for his help with the bioinformatic part in the CAST-Seq pipeline. Chiara Romagnani and Timo Rückert (German Rheumatism Research Center, a Leibniz Institute, Berlin, Germany) for their expert advice on NK cells.

This project has received funding from the European Union’s Horizon 2020 research and innovation program under grant agreement no. 825392 (ReSHAPE-h2020.eu) to M.S.-H., H.-D.V., P.R., and D.L.W.. Further, the project received funding by the European Union under Grant Agreement Nr. 101057438 to T.C., H.-D.V., P.R. and D.L.W.. Views and opinions expressed are however those of the author(s) only and do not necessarily reflect those of the European Union or the European Health and Digital Executive Agency (HADEA). Neither the European Union nor the granting authority can be held responsible for them. Further, J.K. and D.L.W would like to thank the Einstein Center for Regenerative Therapies (ECRT) for support via the ECRT Research Grant (2020-2022) and the ECRT Young Scientist Kickbox grant. Further, J.K. and D.L.W. were supported by the SPARK-BIH program by the Berlin Institute of Health, Germany. M.A. is partially supported by the award No. P30CA014089 from the National Cancer Institute. R.S.’s laboratory was financed by grants of the German Cancer Aid (Deutsche Krebshilfe Nr. 70114234), by The Jackson Laboratory (LV-HLA2) and by a Professorship funded by the Cancer Research Center Cologne Essen (CCCE).

## VII. Author contributions

J.K. designed this study, planned, and performed experiments, analyzed results, interpreted the data, and wrote the manuscript. C.F., V.D., W.D., planned and performed experiments, analyzed results, interpreted the data, and edited the manuscript. V.G., M.St., T.Z., C.P., L.A., J.A. performed experiments and analyzed results. C.F.-G. performed and interpreted CAST-seq and provided respective sections for the manuscript. M.Su., J.H., R.S. planned experiments, interpreted data and edited the manuscript. H.A. provided materials (*Her2-CAR*-transgenes^33^), interpreted the data and edited the manuscript. A.K., M.A. and A.P. provided reagents, interpreted data and edited the manuscript. T.C. supervised work on CAST-seq, provided reagents, interpreted data and edited the manuscript. H.-D.V., P.R., M.S.-H. supervised parts of the study, provided reagents, interpreted data and edited the manuscript. D.L.W. designed and led the study, planned experiments, analyzed results, interpreted data, and wrote the manuscript. All authors reviewed and approved the manuscript in its final form.

## References

1. Kalos M, Levine BL, Porter DL, et al. T cells with chimeric antigen receptors have potent antitumor effects and can establish memory in patients with advanced leukemia. Sci Transl Med. 2011;3(95):95ra73.

2. Liu E, Marin D, Banerjee P, et al. Use of CAR-Transduced Natural Killer Cells in CD19-Positive Lymphoid Tumors. N. Engl. J. Med. 2020;382(6):545–553.

3. MacDonald KG, Hoeppli RE, Huang Q, et al. Alloantigen-specific regulatory T cells generated with a chimeric antigen receptor. J Clin Invest. 2016;126(4):1413–1424.

4. Roemhild A, Otto NM, Moll G, et al. Regulatory T cells for minimising immune suppression in kidney transplantation: phase I/IIa clinical trial. BMJ. 2020;371:.

5. Maude SL, Frey N, Shaw PA, et al. Chimeric antigen receptor T cells for sustained remissions in leukemia. N. Engl. J. Med. 2014;371(16):1507–1517.

6. Neelapu SS, Locke FL, Bartlett NL, et al. Axicabtagene Ciloleucel CAR T-Cell Therapy in Refractory Large B-Cell Lymphoma. N Engl J Med. 2017;377(26):2531–2544.

7. Schuster SJ, Svoboda J, Chong EA, et al. Chimeric Antigen Receptor T Cells in Refractory B-Cell Lymphomas. New England Journal of Medicine. 2017;377(26):2545–2554.

8. Raje N, Berdeja J, Lin Y, et al. Anti-BCMA CAR T-Cell Therapy bb2121 in Relapsed or Refractory Multiple Myeloma. N. Engl. J. Med. 2019;380(18):1726–1737.

9. Geisler C, Kuhlmann J, Rubin B. Assembly, intracellular processing, and expression at the cell surface of the human alpha beta T cell receptor/CD3 complex. Function of the CD3-zeta chain. J Immunol. 1989;143(12):4069–4077.

10. Irving BA, Weiss A. The cytoplasmic domain of the T cell receptor zeta chain is sufficient to couple to receptor-associated signal transduction pathways. Cell. 1991;64(5):891–901.

11. Moingeon P, Lucich JL, McConkey DJ, et al. CD3 zeta dependence of the CD2 pathway of activation in T lymphocytes and natural killer cells. PNAS. 1992;89(4):1492–1496.

12. Omer B, Cardenas MG, Pfeiffer T, et al. A Costimulatory CAR Improves TCR-based Cancer Immunotherapy. Cancer Immunology Research. 2022;10(4):512–524.

13. Majzner RG, Ramakrishna S, Yeom KW, et al. GD2-CAR T cell therapy for H3K27M-mutated diffuse midline gliomas. Nature. 2022;603(7903):934–941.

14. Mougiakakos D, Krönke G, Völkl S, et al. CD19-Targeted CAR T Cells in Refractory Systemic Lupus Erythematosus. New England Journal of Medicine. 2021;385(6):567–569.

15. Mackensen A, Müller F, Mougiakakos D, et al. Anti-CD19 CAR T cell therapy for refractory systemic lupus erythematosus. Nat Med. 2022;28(10):2124–2132.

16. Müller F, Boeltz S, Knitza J, et al. CD19-targeted CAR T cells in refractory antisynthetase syndrome. The Lancet. 2023;0(0):

17. Eyquem J, Mansilla-Soto J, Giavridis T, et al. Targeting a CAR to the TRAC locus with CRISPR/Cas9 enhances tumour rejection. Nature. 2017;543(7643):113–117.

18. Odak A, Yuan H, Feucht J, et al. Novel extragenic genomic safe harbors for precise therapeutic T cell engineering. Blood. 2023;blood.2022018924.

19. Scholler J, Brady TL, Binder-Scholl G, et al. Decade-Long Safety and Function of Retroviral-Modified Chimeric Antigen Receptor T Cells. Science Translational Medicine. 2012;4(132):132ra53-132ra53.

20. Fraietta JA, Nobles CL, Sammons MA, et al. Disruption of TET2 promotes the therapeutic efficacy of CD19-targeted T cells. Nature. 2018;558(7709):307–312.

21. Shah NN, Qin H, Yates B, et al. Clonal expansion of CAR T cells harboring lentivector integration in the CBL gene following anti-CD22 CAR T-cell therapy. Blood Adv. 2019;3(15):2317–2322.

22. Micklethwaite KP, Gowrishankar K, Gloss BS, et al. Investigation of product-derived lymphoma following infusion of piggyBac-modified CD19 chimeric antigen receptor T cells. Blood. 2021;138(16):1391–1405.

23. Bishop DC, Clancy LE, Simms R, et al. Development of CAR T-cell lymphoma in 2 of 10 patients effectively treated with piggyBac-modified CD19 CAR T cells. Blood. 2021;138(16):1504–1509.

24. Wagner DL, Koehl U, Chmielewski M, Scheid C, Stripecke R. Review: Sustainable Clinical Development of CAR-T Cells - Switching From Viral Transduction Towards CRISPR-Cas Gene Editing. Front Immunol. 2022;13:865424.

25. Müller TR, Jarosch S, Hammel M, et al. Targeted T cell receptor gene editing provides predictable T cell product function for immunotherapy. Cell Rep Med. 2021;2(8):100374.

26. MacLeod DT, Antony J, Martin AJ, et al. Integration of a CD19 CAR into the TCR Alpha Chain Locus Streamlines Production of Allogeneic Gene-Edited CAR T Cells. Molecular Therapy. 2017;25(4):949–961.

27. Dai X, Park JJ, Du Y, et al. One-step generation of modular CAR-T cells with AAV-Cpf1. Nat Methods. 2019;16(3):247–254.

28. Wiebking V, Lee CM, Mostrel N, et al. Genome editing of donor-derived T-cells to generate allogenic chimeric antigen receptor-modified T cells: Optimizing αβ T cell-depleted haploidentical hematopoietic stem cell transplantation. Haematologica. 2020;haematol.2019.233882.

29. Zhang J, Hu Y, Yang J, et al. Non-viral, specifically targeted CAR-T cells achieve high safety and efficacy in B-NHL. Nature. 2022;609(7926):369–374.

30. Allen AG, Khan SQ, Margulies CM, et al. A highly efficient transgene knock-in technology in clinically relevant cell types. Nat Biotechnol. 2023;

31. Torikai H, Reik A, Liu P-Q, et al. A foundation for universal T-cell based immunotherapy: T cells engineered to express a CD19-specific chimeric-antigen-receptor and eliminate expression of endogenous TCR. Blood. 2012;119(24):5697–5705.

32. Kath J, Du W, Pruene A, et al. Pharmacological interventions enhance virus-free generation of TRAC-replaced CAR T cells. Molecular Therapy - Methods & Clinical Development. 2022;25:311–330.

33. Nguyen DN, Roth TL, Li PJ, et al. Polymer-stabilized Cas9 nanoparticles and modified repair templates increase genome editing efficiency. Nat Biotechnol. 2020;38(1):44–49.

34. Turchiano G, Andrieux G, Klermund J, et al. Quantitative evaluation of chromosomal rearrangements in gene-edited human stem cells by CAST-Seq. Cell Stem Cell. 2021;28(6):1136–1147.e5.

35. Rhiel M, Geiger K, Andrieux G, et al. T-CAST: An optimized CAST-Seq pipeline for TALEN confirms superior safety and efficacy of obligate-heterodimeric scaffolds. Front Genome Ed. 2023;5:1130736.

36. Textor A, Listopad JJ, Wührmann LL, et al. Efficacy of CAR T-cell therapy in large tumors relies upon stromal targeting by IFNγ. Cancer Res. 2014;74(23):6796–6805.

37. Hermans IF, Silk JD, Yang J, et al. The VITAL assay: a versatile fluorometric technique for assessing CTL- and NKT-mediated cytotoxicity against multiple targets in vitro and in vivo. Journal of Immunological Methods. 2004;285(1):25–40.

38. Künkele A, Taraseviciute A, Finn LS, et al. Preclinical Assessment of CD171-Directed CAR T-cell Adoptive Therapy for Childhood Neuroblastoma: CE7 Epitope Target Safety and Product Manufacturing Feasibility. Clin Cancer Res. 2017;23(2):466–477.

39. Yang S, Cohen CJ, Peng PD, et al. Development of optimal bicistronic lentiviral vectors facilitates high-level TCR gene expression and robust tumor cell recognition. Gene therapy. 2008;15(21):1411.

40. Liu Z, Chen O, Wall JBJ, et al. Systematic comparison of 2A peptides for cloning multi-genes in a polycistronic vector. Sci Rep. 2017;7(1):2193.

41. Ahmed N, Brawley VS, Hegde M, et al. Human Epidermal Growth Factor Receptor 2 (HER2) -Specific Chimeric Antigen Receptor-Modified T Cells for the Immunotherapy of HER2-Positive Sarcoma. J. Clin. Oncol. 2015;33(15):1688–1696.

42. Ahmed N, Brawley V, Hegde M, et al. HER2-Specific Chimeric Antigen Receptor–Modified Virus-Specific T Cells for Progressive Glioblastoma. JAMA Oncol. 2017;3(8):1094–1101.

43. Hegde M, Joseph SK, Pashankar F, et al. Tumor response and endogenous immune reactivity after administration of HER2 CAR T cells in a child with metastatic rhabdomyosarcoma. Nature Communications. 2020;11(1):3549.

44. Noyan F, Zimmermann K, Hardtke-Wolenski M, et al. Prevention of Allograft Rejection by Use of Regulatory T Cells With an MHC-Specific Chimeric Antigen Receptor. American Journal of Transplantation. 2017;17(4):917–930.

45. Deniger DC, Switzer K, Mi T, et al. Bispecific T-cells expressing polyclonal repertoire of endogenous γδ T-cell receptors and introduced CD19-specific chimeric antigen receptor. Mol Ther. 2013;21(3):638–647.

46. Daher M, Rezvani K. Outlook for New CAR-Based Therapies with a Focus on CAR NK Cells: What Lies Beyond CAR-Engineered T Cells in the Race against Cancer. Cancer Discovery. 2021;11(1):45–58.

47. Klingemann H. The NK-92 cell line-30 years later: its impact on natural killer cell research and treatment of cancer. Cytotherapy. 2023;25(5):451–457.

48. Anderson P, Caligiuri M, O’Brien C, et al. Fc gamma receptor type III (CD16) is included in the zeta NK receptor complex expressed by human natural killer cells. Proceedings of the National Academy of Sciences. 1990;87(6):2274–2278.

49. Gong JH, Maki G, Klingemann HG. Characterization of a human cell line (NK-92) with phenotypical and functional characteristics of activated natural killer cells. Leukemia. 1994;8(4):652–658.

50. Qasim W, Zhan H, Samarasinghe S, et al. Molecular remission of infant B-ALL after infusion of universal TALEN gene-edited CAR T cells. Sci Transl Med. 2017;9(374):.

51. Benjamin R, Graham C, Yallop D, et al. Genome-edited, donor-derived allogeneic anti-CD19 chimeric antigen receptor T cells in paediatric and adult B-cell acute lymphoblastic leukaemia: results of two phase 1 studies. The Lancet. 2020;396(10266):1885–1894.

52. Qasim W. Genome edited allogeneic donor “universal” chimeric antigen receptor T Cells. Blood. 2022;blood.2022016204.

53. Wagner DL, Fritsche E, Pulsipher MA, et al. Immunogenicity of CAR T cells in cancer therapy. Nature Reviews Clinical Oncology. 2021;1–15.

54. Prinzing B, Zebley CC, Petersen CT, et al. Deleting DNMT3A in CAR T cells prevents exhaustion and enhances antitumor activity. Sci Transl Med. 2021;13(620):eabh0272.

55. Carnevale J, Shifrut E, Kale N, et al. RASA2 ablation in T cells boosts antigen sensitivity and long-term function. Nature. 2022;609(7925):174–182.

56. Wiebking V, Patterson JO, Martin R, et al. Metabolic engineering generates a transgene-free safety switch for cell therapy. Nature Biotechnology. 2020;1–10.

57. Torikai H, Reik A, Soldner F, et al. Toward eliminating HLA class I expression to generate universal cells from allogeneic donors. Blood. 2013;122(8):1341–1349.

58. Kagoya Y, Guo T, Yeung B, et al. Genetic Ablation of HLA Class I, Class II, and the T-cell Receptor Enables Allogeneic T Cells to Be Used for Adoptive T-cell Therapy. Cancer Immunol Res. 2020;8(7):926–936.

59. Pomeroy EJ, Hunzeker JT, Kluesner MG, et al. A Genetically Engineered Primary Human Natural Killer Cell Platform for Cancer Immunotherapy. Mol Ther. 2020;28(1):52–63.

60. Daher M, Basar R, Gokdemir E, et al. Targeting a cytokine checkpoint enhances the fitness of armored cord blood CAR-NK cells. Blood. 2021;137(5):624–636.

61. Diorio C, Murray R, Naniong M, et al. Cytosine Base Editing Enables Quadruple-Edited Allogeneic CAR-T Cells for T-ALL. Blood. 2022;

62. Komor AC, Kim YB, Packer MS, Zuris JA, Liu DR. Programmable editing of a target base in genomic DNA without double-stranded DNA cleavage. Nature. 2016;533(7603):420–424.

63. Gaudelli NM, Komor AC, Rees HA, et al. Programmable base editing of A•T to G•C in genomic DNA without DNA cleavage. Nature. 2017;551(7681):464–471.

64. Glaser V, Flugel C, Kath J, et al. Combining different CRISPR nucleases for simultaneous knock-in and base editing prevents translocations in multiplex-edited CAR T cells. Genome Biology. 2023;24(1):89.

65. Shy BR, Vykunta VS, Ha A, et al. High-yield genome engineering in primary cells using a hybrid ssDNA repair template and small-molecule cocktails. Nat Biotechnol. 2022;

66. Barden M, Holzinger A, Velas L, et al. CAR and TCR form individual signaling synapses and do not cross-activate, however, can co-operate in T cell activation. Front Immunol. 2023;14:1110482.

67. Kath J, Du W, Martini S, et al. CAR NK-92 cell-mediated depletion of residual TCR+ cells for ultrapure allogeneic TCR-deleted CAR T-cell products. Blood Adv. 2023;7(15):4124–4134.

68. Weber EW, Parker KR, Sotillo E, et al. Transient rest restores functionality in exhausted CAR-T cells through epigenetic remodeling. Science. 2021;372(6537):eaba1786.

69. Ruella M, Xu J, Barrett DM, et al. Induction of resistance to chimeric antigen receptor T cell therapy by transduction of a single leukemic B cell. Nature Medicine. 2018;24(10):1499–1503.

70. Roberts JL, Lauritsen JPH, Cooney M, et al. T−B+NK+ severe combined immunodeficiency caused by complete deficiency of the CD3ζ subunit of the T-cell antigen receptor complex. Blood. 2007;109(8):3198–3206.

71. Valés-Gómez M, Esteso G, Aydogmus C, et al. Natural killer cell hyporesponsiveness and impaired development in a CD247-deficient patient. Journal of Allergy and Clinical Immunology. 2016;137(3):942–945.e4.

72. Dahlvang JD, Dick JK, Sangala JA, et al. Ablation of SYK Kinase from Expanded Primary Human NK Cells via CRISPR/Cas9 Enhances Cytotoxicity and Cytokine Production. J Immunol. 2023;ji2200488.

73. Attaf M, Legut M, Cole DK, Sewell AK. The T cell antigen receptor: the Swiss army knife of the immune system. Clinical and Experimental Immunology. 2015;181(1):1–18.

74. Gomes-Silva D, Mukherjee M, Srinivasan M, et al. Tonic 4-1BB Costimulation in Chimeric Antigen Receptors Impedes T Cell Survival and Is Vector Dependent. Cell Rep. 2017;21(1):17–26.

75. Rodriguez-Marquez P, Calleja-Cervantes ME, Serrano G, et al. CAR density influences antitumoral efficacy of BCMA CAR T cells and correlates with clinical outcome. Sci Adv. 2022;8(39):eabo0514.

76. Ho J-Y, Wang L, Liu Y, et al. Promoter usage regulating the surface density of CAR molecules may modulate the kinetics of CAR-T cells in vivo. Molecular Therapy - Methods & Clinical Development. 2021;21:237–246.

77. Flugel CL, Majzner RG, Krenciute G, et al. Overcoming on-target, off-tumour toxicity of CAR T cell therapy for solid tumours. Nat Rev Clin Oncol. 2023;20(1):49–62.

