## Supplementary Figures for "Integration of ζ-deficient CARs into the *CD3-zeta* gene conveys potent cytotoxicity in T and NK cells"

### Supplementary Figures – Content overview:


**Page**

- Supplementary Figure 1: Acute lymphoblastic leukemia xenograft mouse model using luciferase-labeled Nalm-6 ($CD19^+$) tumor cells. 2
    - *Additional data supporting Figure 1.*
- Supplementary Figure 2: Tuning of CAR-expression from *CD3ζ* gene increases CAR-dependent production of cytokines. 3
    - *Additional data supporting Figure 2.*
- Supplementary Figure 3: Tuning of CAR-expression from *CD3ζ* gene does not significantly influence exhaustion status. 4
    - *Additional data supporting Figure 2.*
- Supplementary Figure 4: After repetitive co-culture, *TRAC* and *CD3ζ* integrated CAR T cells have a similar phenotype and show preserved differences in CAR-dependent cytokine expression. 5
    - *Additional data supporting Figure 2.*
- Supplementary Figure 5: Extended CAR T cells expansion results in more differentiated, $T_{EM}$ dominated phenotype. 6
    - *Additional Data supporting Figure 2.*
- Supplementary Figure 6: *CD3ζ truncCAR* knock-in generates HER2-specific CAR T cells that display less signs of differentiation and reduced activation induced cell death. 7
- Supplementary Figure 7: Off-target analysis for *CD3ζ*-targeting sgRNA using CAST-Seq. 8
- Supplementary Figure 8: Summary for editing efficiency in other non-conventional T cell types. 9
    - *Additional data supporting Figure 3.*
- Supplementary Figure 9: *CD3ζ*-disuption does not impede canonical NK cell functions *in vitro*. 10
    - *Additional data supporting Figure 4.*
- Supplementary Figure 10: Analysis of CD16 expression on different NK cells and evaluation of primary NK cytotoxicity toward Jeko-1 cell line. 11
    - *Additional data supporting Figure 4.*

**Supplementary Figure 1: Acute lymphoblastic leukemia xenograft mouse model using luciferase-labeled Nalm-6 (CD19⁺) tumor cells.**

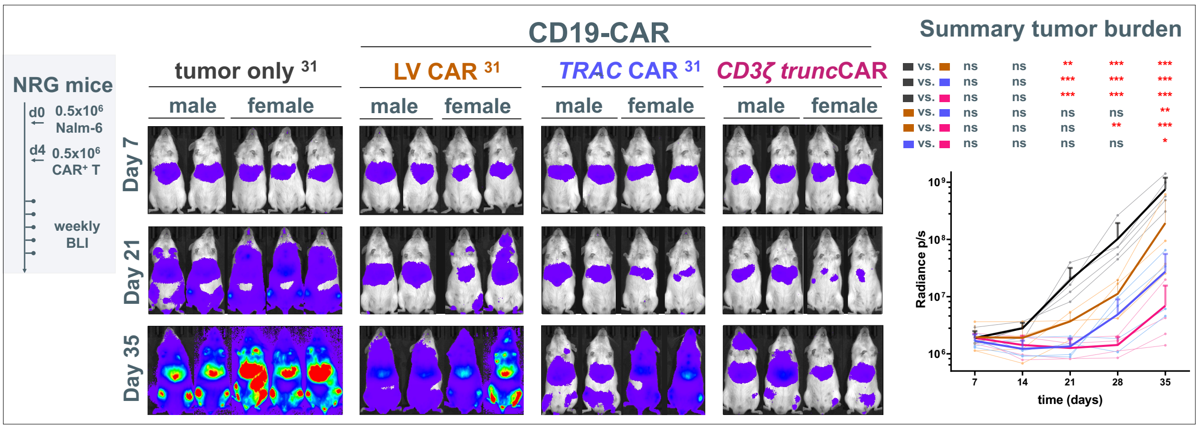


**Suppl. Fig. 1: Acute lymphoblastic leukemia xenograft mouse model using luciferase-labeled Nalm-6 (CD19$^+$) tumor cells**. 4 days post administration of 0.5x10$^6$ luciferase-expressing Nalm-6 cells, fresh, 14-day expanded, TCR-deleted CAR T cells were injected systemically at a dose of 0.5x10$^6$ CAR$^+$ cells/mouse. Tumor burden was assessed via bioluminescence imaging (BLI). The control groups in animal experiment 2 were previously published in Kath *et al.* (2022) (Ref. 31 in main manuscript). Statistics: n=4-5; BLI data were log-transformed and compared using a repeated measures 2-way ANOVA with Geisser-Greenhouse correction followed by a Dunnett's multiple comparison test.

**Supplementary Figure 2: Tuning of CAR-expression from *CD3ζ* gene increases CAR-dependent production of cytokines.**

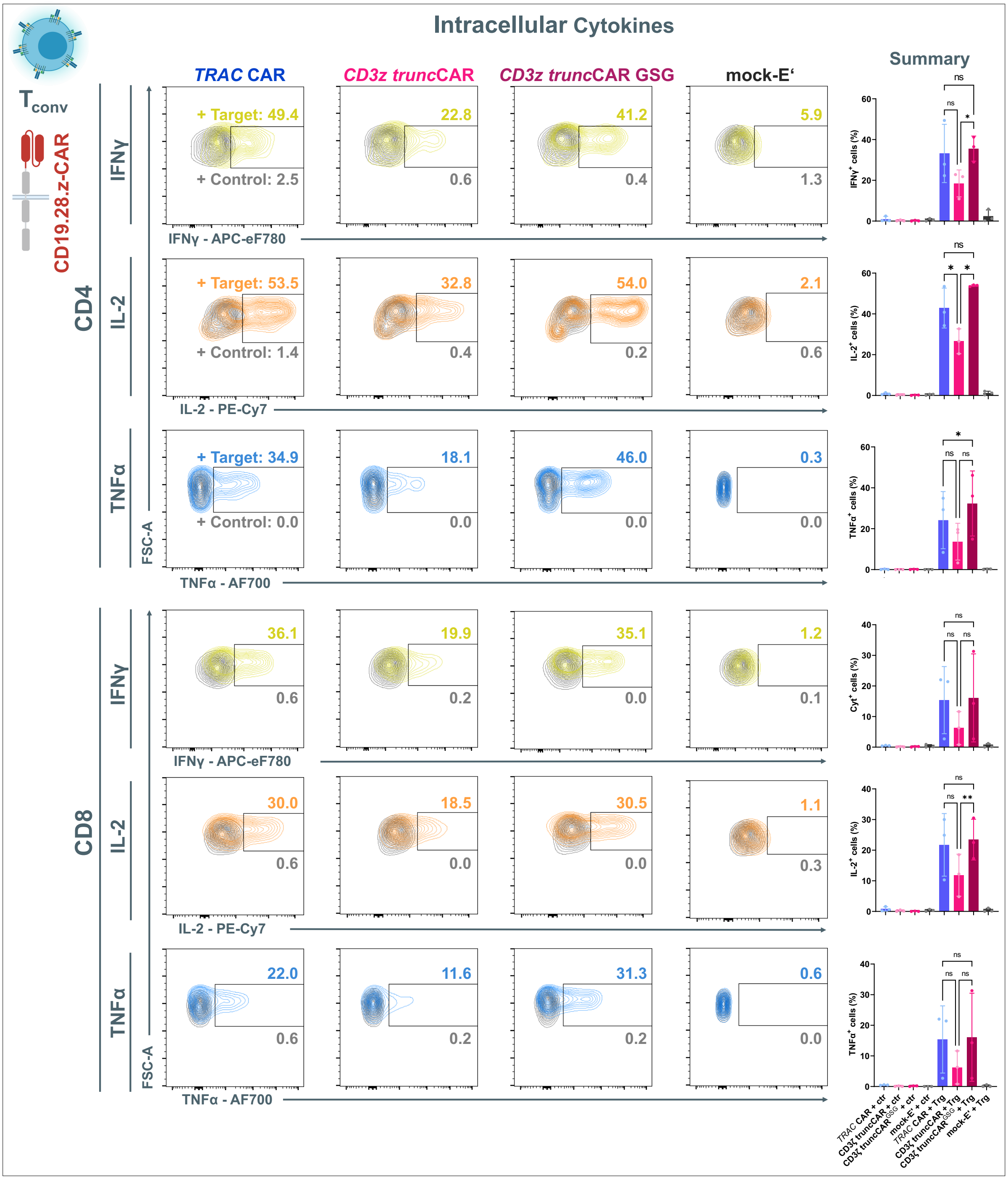


**Suppl. Fig. 2: Tuning of CAR-expression from *CD3ζ* gene increases CAR-dependent production of cytokines.** Intracellular detection of cytokines in CAR$^+$ T cells via flow cytometry after encounter with target (Trg) cells (colored graphs in contour plots) or control (ctr) cells (grey plots). Summary plots on the right: n=3 biol. repl.; statistics: repeated measures one-way ANOVA followed by uncorrected Fischer's LSD, with a single pooled variance.

**Supplementary Figure 3: Tuning of CAR-expression from *CD3ζ* gene does not significantly influence exhaustion status.**

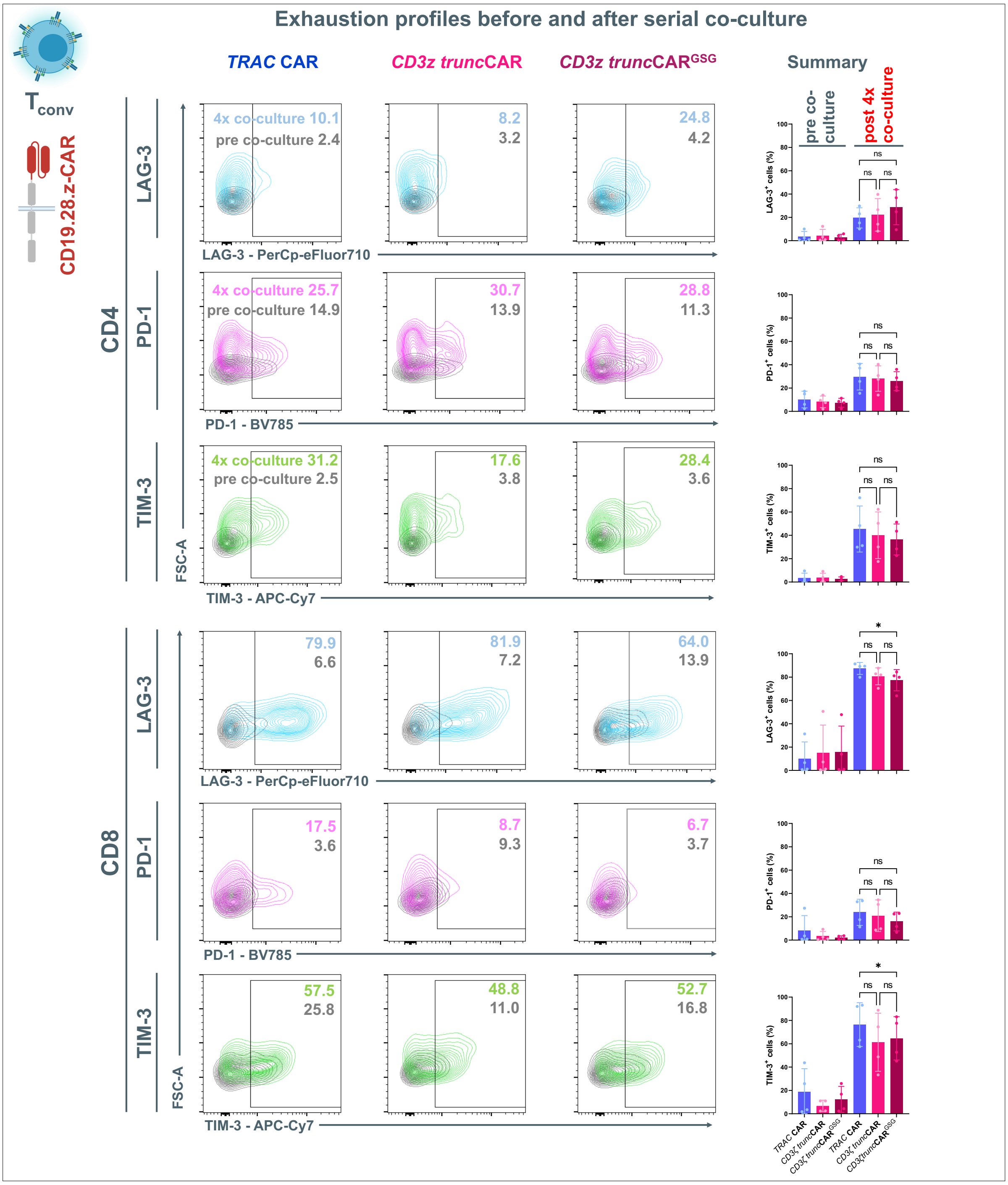


**Suppl. Fig. 3: Tuning of CAR-expression from *CD3ζ* gene does not significantly influence exhaustion status.** Flow cytometric detection of the inhibitory surface receptors LAG-3, PD-1 and TIM-3 on CAR⁺ T cells before (grey graphs in contour plots) or after repetitive co-culture (colored plots). Summary plots on the right: n=4 biol. repl.; statistics: repeated measures one-way ANOVA followed by uncorrected Fischer's LSD, with a single pooled variance.

**Supplementary Figure 4: After repetitive co-culture, *TRAC* and *CD3ζ* integrated CAR T cells have a similar phenotype and show preserved differences in CAR-dependent cytokine expression.**

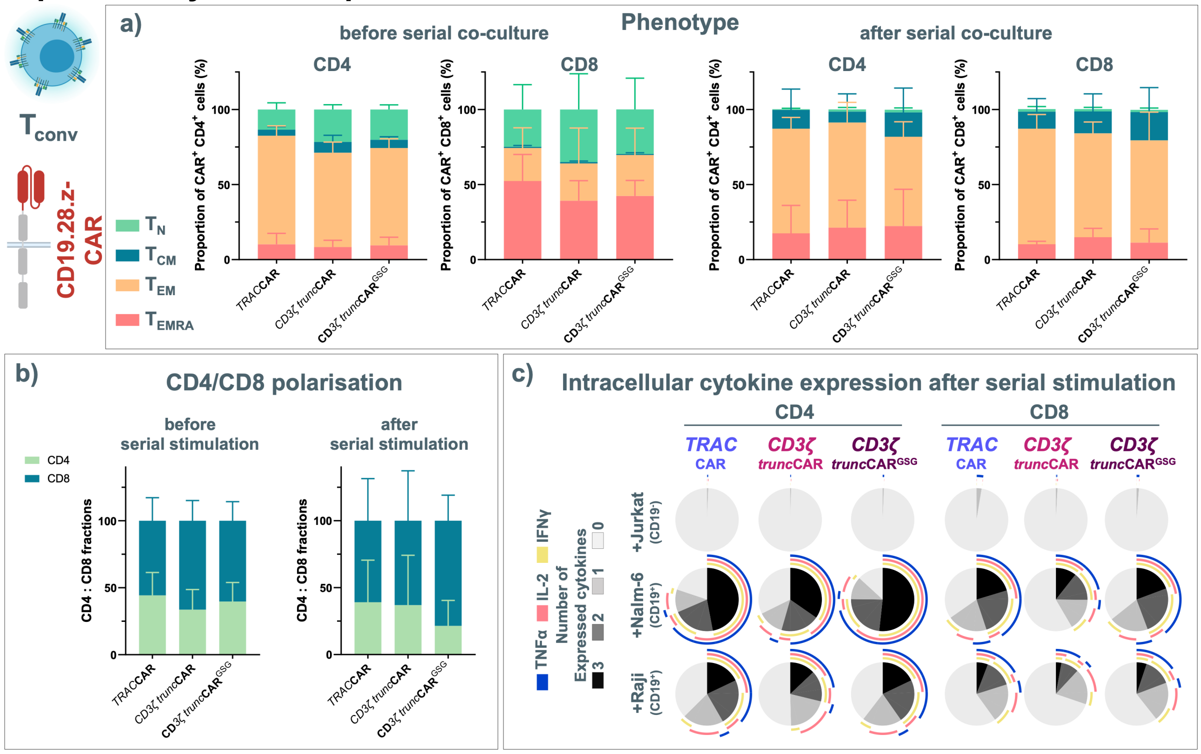


**Suppl. Fig. 4: After repetitive co-culture, *TRAC* and *CD3ζ* integrated CAR T cells have a similar phenotype and show preserved differences in CAR-dependent cytokine expression.** (**a**) Phenotype characterization of CAR⁺ T cells before serial Nalm-6 co-culture (day 14 post blood collection) and after co-culture via flow cytometric detection of CCR7 and CD45RA ($T_N$ (naïve-like): CCR7$^+$ CD45RA$^+$; $T_{CM}$ (central memory): CCR7$^+$ CD45RA$^-$; $T_{EM}$ (effector memory): CCR7$^-$ CD45RA$^-$; $T_{EMRA}$ (terminally differentiated effector memory RA$^+$): CCR7$^-$ CD45RA$^+$); (n=4 biol. repl.); (**b**) CD4/CD8 polarization detected via flow cytometry before and after serial Nalm-6 cell co-culture (n=4 biol. repl.); (**c**) intracellularly detected cytokines after serial Nalm-6 cell co-culture (n=4 biol. repl.)

**Supplementary Figure 5: Extended CAR T cells expansion results in more differentiated, $T_{EM}$ dominated phenotype.**

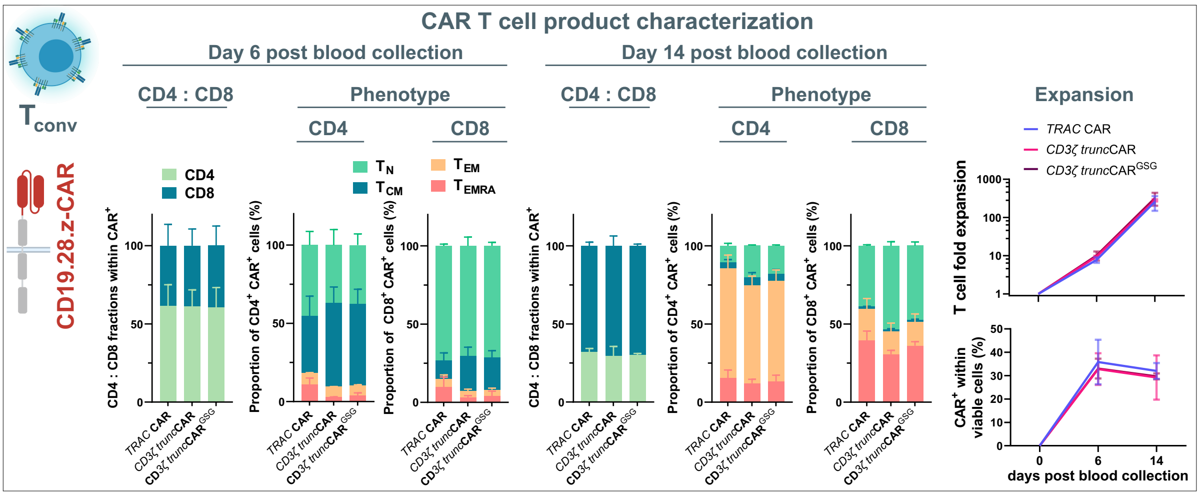


**Suppl. Fig. 5: Extended CAR T cells expansion results in more differentiated, $T_{EM}$ dominated phenotype.** Left: Phenotype characterization of G-rex-expanded CAR+ cells on day 6 and day 14 post blood collection. Right: T cell fold expansion and CAR+ frequency during expansion. (n=2 biol. repl.)

**Supplementary Figure 6: *CD3ζ trunc*CAR knock-in generates HER2-specific CAR T cells that display less signs of differentiation and reduced activation induced cell death.**

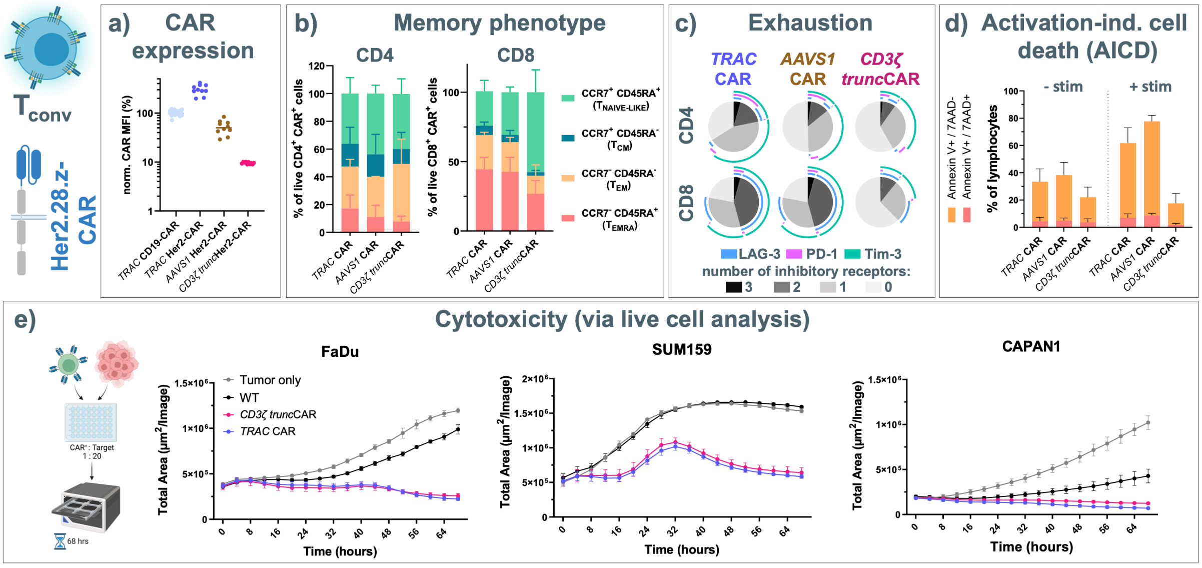


**Suppl. Fig. 6: *CD3ζ trunc*CAR knock-in generates HER2-specific CAR T cells that display less signs of differentiation and reduced activation induced cell death.** (**a**) Mean fluorescence intensity (MFI) determined by flow cytometry as a measure of cellular CAR expression normalized to the *TRAC*-CD19-CAR condition (n=2 biol. repl., each in 5 techn. repl.); (**b**) CAR$^{+}$ T cell phenotype (n=3 biol. repl.); (**c**) Cell surface expression of inhibitory receptors as a measure of cellular exhaustion (n=3 biol. repl.); (**d**) Activation-induced cell death assessed by flow cytometry via staining of Annexin V and 7AAD. (n=2 donors, each in 2 techn. repl.) (**e**) *In vitro* tumor control assessed via live cell imaging (CAR+ T cell : Tumor cell ratio of 1:20. Mean +/- S.D., n=4 techn. repl.).

**Supplementary Figure 7. Off-target analysis for *CD3ζ*-targeting sgRNA using CAST-Seq.**

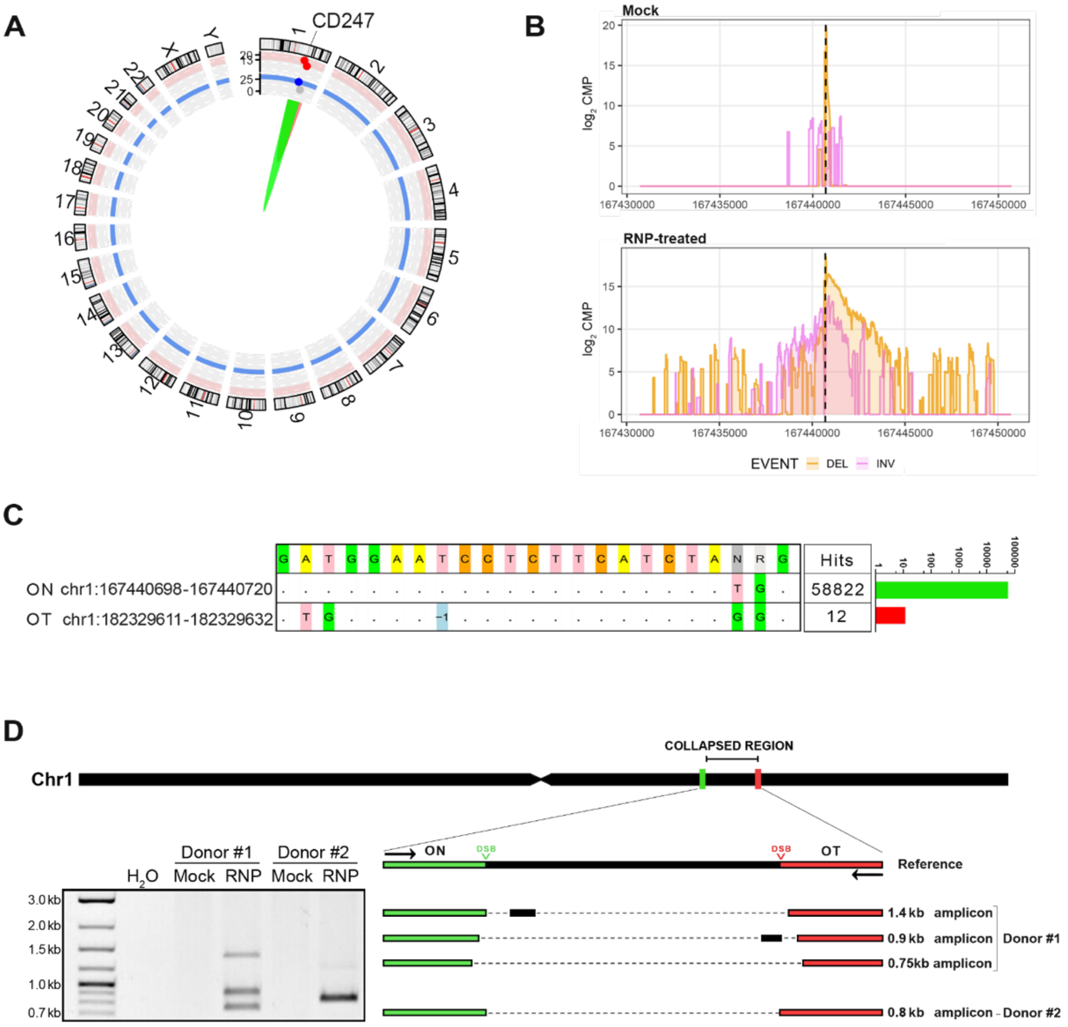


**Suppl. Fig. 7. Off-target analysis for *CD3ζ*-targeting sgRNA using CAST-Seq.** (**A**) Structural variations. Circos plot shows results of chromosomal rearrangements detected in cells edited with RNPs. On-target (ON) genomic aberrations are indicated in green, an off-target mediated rearrangement between the *CD247* target site and an off-target (OT) site on the same chromosome in red. (**B**) On-target aberrations. CAST-Seq coverage plots show reads aligned to the +/-10 kb regions flanking the *CD247* target site. Genomic aberrations in form of deletions (DEL) and inversions (INV) are shown in orange and purple, respectively. The x-axis represents the chromosomal coordinates, the y-axis the log2 read count per million (CPM), and the dotted line the cleavage site. (**C**) Off-target site. Alignment of the nominated OT site to the *CD247* target site and number of CAST-Seq hits. (**D**) Validation of off-target activity. Schematic on top shows locations of on- and off-target sites on chromosome 1 and primer binding sites. Result of PCR analysis is shown on the left, schematic representing the Sanger sequencing results on the right. The analysis was performed in cells from two different donors.

**Supplementary Figure 8: Summary for editing efficiency in other non-conventional T cell types.**

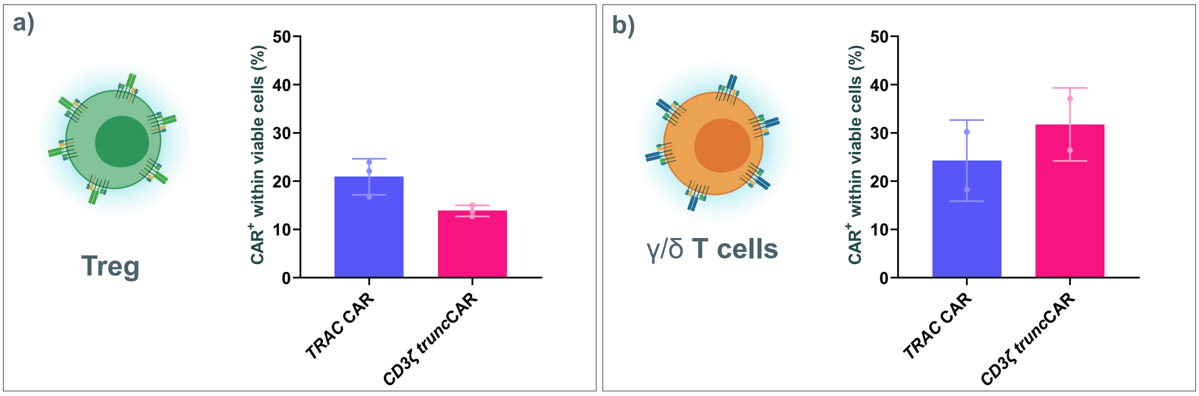


**Suppl. Figure 8: Summary for editing efficiency in other non-conventional T cell types.** (**a**) $T_{reg}$ n=3 biol. repl. (**b**) $TCR_{\gamma/\delta}$ T cells. n=2 biol. repl.

**Supplementary Figure 9: *CD3ζ*-disuption does not impede canonical NK cell functions *in vitro*.**

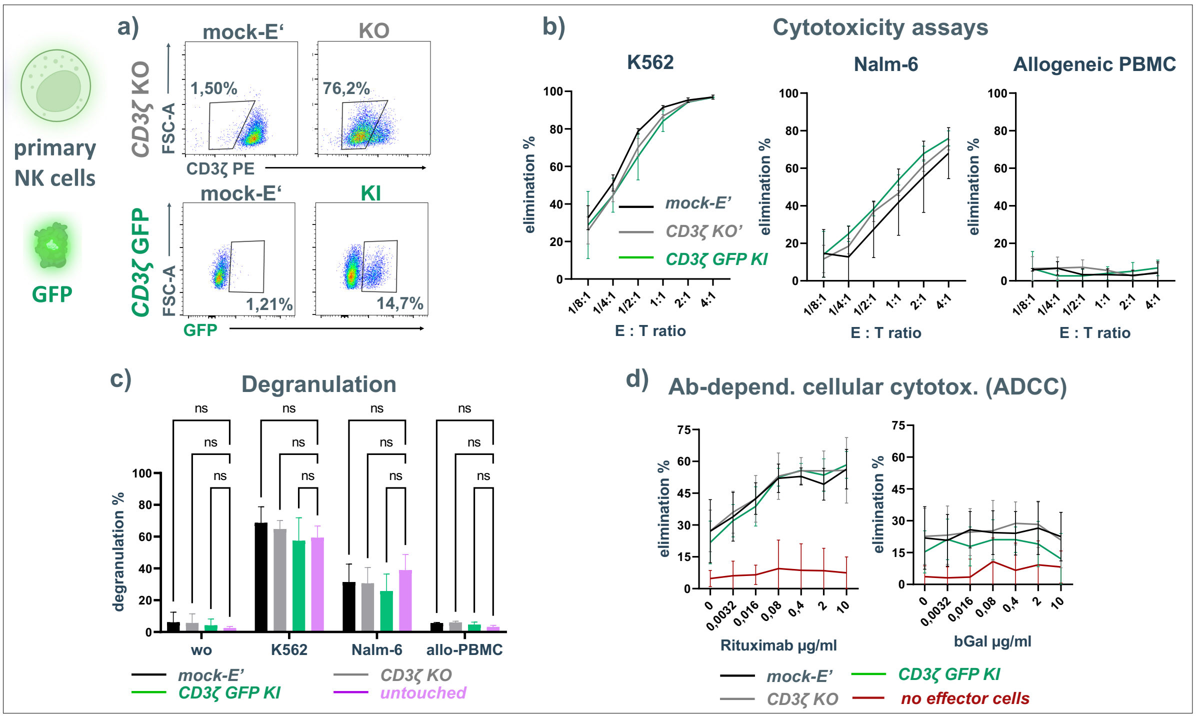


**Suppl. Fig. 9: *CD3ζ*-disuption does not impede canonical NK cell functions *in vitro*.** (**a**) *CD3ζ* editing outcomes assessed by flow cytometry. (**b**) Cytotoxicity of primary *CD3ζ* disrupted NK cells in simple 16h co-culture assay with K562 cells, Nalm-6 cells or allogeneic PBMC. (n=3 biol. repl.); (**c**) Degranulation of primary *CD3ζ* disrupted NK cells assessed by flow cytometry. (n = 3 biol. repl.; two-way ANOVA followed by Dunnett's multiple comparison test with a single pooled variance); (**d**) ADCC of primary *CD3ζ* disrupted NK cells against $CD20^+$ $bGal^-$ Jeko-1 cells at different concentrations of antibodies specific for CD20 (Rituximab) or bGal (n=3 biol. repl., each in 3 techn. repl.).

**Supplementary Figure 10: Analysis of CD16 expression on different NK cells and evaluation of primary NK cytotoxicity toward Jeko-1 cell line.**

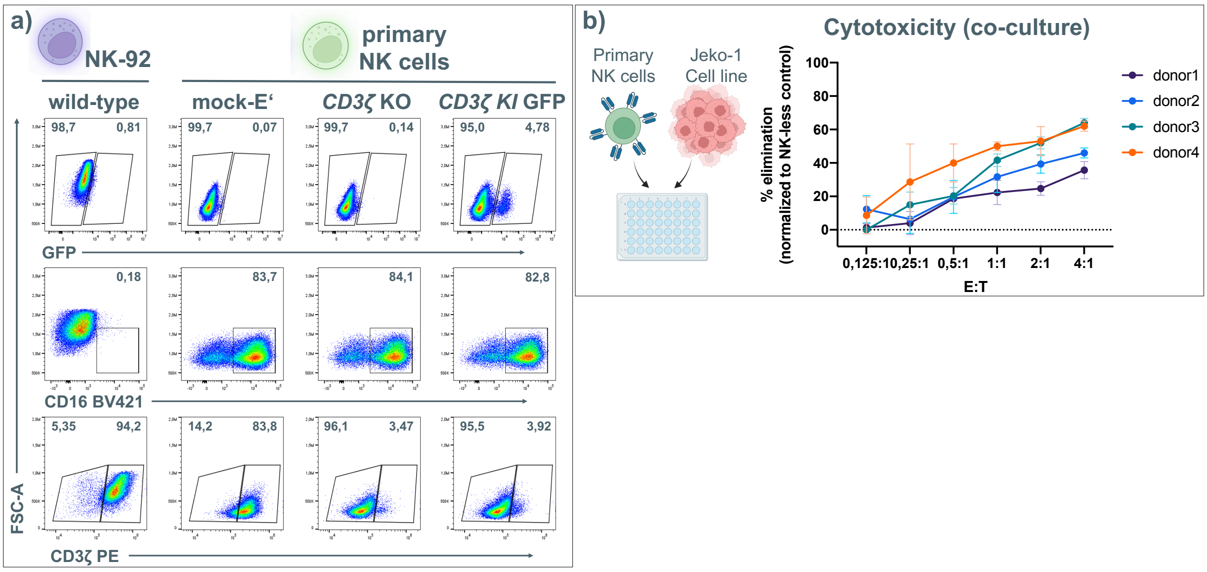


**Suppl. Fig. 10: Analysis of CD16 expression on different NK cells and evaluation of primary NK cytotoxicity toward Jeko-1 cell line.** (a) Expression of CD16 and *CD3ζ* in wild-type NK-92 cells and primary NK cells after *CD3ζ*-disruption. (**b**) NK mediated cytotoxicity against Jeko-1 cells in *vitro*. (n = 4 biol. repl., each in 3 techn. repl.)
