## Supplementary Methods for "Integration of ζ-deficient CARs into the *CD3-zeta* gene conveys potent cytotoxicity in T and NK cells"

*Tumor cell line culture*

NK-92 cells were cultured in RPMI supplemented with 10% FCS and IL-2 (500 IU/ml). Nalm-6- , Raji-, Jeko-1-derived cell lines were cultured in RPMI medium supplemented with 10% FCS. GFP- and luciferase-expressing Fadu, Sum159 and Capan1 cells were generated by Suzuki Laboratory (BCM) and cultured in IMDM medium supplemented with 10% FCS and 1% penicillin/streptomycin.

*Cytotoxicity assessment*

CAR-independent cytotoxicity of NK-cells was assessed in 4-hour (primary NK cells) or 16-hour (NK-92 cells) co-cultures with the respective target cell line at a range of Effector cell (E) to Target cell (T) ratios. Killing was assessed via cell counts of surviving target cells. CAR-dependent cytotoxicity of CD19-CAR-engineered T and NK cells was assessed in VITAL assays^1^, which are co-culture assays with three cell types: 1.) the effector cells, here CD19-CAR T or NK cells; 2.) target cells, here GFP^+^ Nalm-6 cells (CD19^+^); and 3.) control cells (C), here RFP^+^ CD19-KO Nalm-6 cells. After 4h (for CAR NK cells), 6h (for CAR T cells) of co-culture, ratios of surviving target and control cells were assessed. Relative cytotoxicity was normalized against control wells without effector cells using the formula:

*Relative Cytotoxicity* *= 1 – ([T:C]_sample_/[T:C]_control_)*

*Intracellular cytokine detection*

In separate wells on a round-bottom 96-well plate, CD19-CAR T cells were co-cultured either with CD19^-^ Jurkat cells serving as control cells or with one of the CD19^+^ target cell lines Nalm-6 and Raji. Prior to the assay, target and control cells were labeled with carboxyfluorescein diacetate succinimidyl ester (CFDA-SE, Thermo Fisher Scientific). 1 hour after the start of the co-culture, Brefeldin A (Sigma Aldrich) was added (at 10 μg/mL). After 16 hours, cells were washed in PBS and stained with fixable blue dead cell stain dye for discrimination of dead cells. After another washing step, the cells were fixed and permeabilized using the Intracellular Fixation & Permeabilization Buffer Set (Thermo Fisher Scientific) prior to intracellular staining (**Suppl. Table 2**).

*Nalm-6 rechallenge and exhaustion panel*

T cells were seeded into 24-well plates at a density of 150.000 CD19-CAR^+^ cells/well and co-cultured with 150.000 Nalm-6 target cells in a 1:1 mixture of RPMI and CTL media supplemented with 10% FCS and IL-2 (50 IU/ml). Every 7 days, CAR T cells were re-adjusted to 150.000 CAR^+^ cells/well prior to the next round of co-cultures. After the 4th co-culture, CAR T cells were analyzed by flow cytometry for expression of the inhibitory receptors PD-1, LAG-3 and Tim-3

Mouse study 1 (1x10^6^ CAR^+^ T cell dose)

Mouse study 1 was performed by EPO Berlin GmbH in Berlin, Germany. Animal studies were performed in accordance with the German Animal Welfare Act and approved by local authorities (Landesamt für Gesundheit und Soziales, LaGeSo Berlin, Germany) under the permission A0010/19 for in vivo therapy experiments. Eight-ten-week-old NOD/Shi^-scid^/IL-2Rγ^null^ (NOG) mice were infused with 0.5x10^6^ Nalm-6 cells (expressing GFP and firefly luciferase) via tail vein injection. Four days later, 1x10^6^ TCR-deficient CD19-CAR T cells were infused intravenously. Those CAR T cells were generated either via targeted integration of a CAR or a t*runc*CAR into the TRAC or CD3ζ gene, respectively, or by LV gene transfer and consecutive *TRAC*-KO. All cell products were depleted of residual TCR/CD3^+^ T cells via MACS (CD3 microbeads and LD columns, Miltenyi). CAR T cells were cry-preserved in FCS supplemented with 10% DMSO on day 14 post blood collection and thawed and formulated in PBS 3 hours prior to application. Tumor burden was assessed using non-invasive bioluminescence imaging (BLI). Mice were anesthetized with Isoflurane (Baxter, San Juan, Puerto Rico) and received intraperitoneally 150 mg/kg D-Luciferin (Biosynth, Staad, Switzerland) dissolved in PBS. BLI was performed with the NightOwl II LB983 in vivo imaging system. It was installed with an anaesthesia option for Isoflurane. For device control and initial data analysis the software IndiGO 2.0.5.0 is used. ImageJ software version 1.50i was used for quantification and color-coding of the signal intensity. Overlay pictures were created with Adobe Photoshop CS5.1 software. The animal welfare was controlled twice daily. Body weights and general health conditions were recorded throughout the whole study. Mice were euthanized after individual evaluation for each mouse with BWL >20% or ethical end points reached (health score < 2). After 60 days all remaining mice: final necropsy for leukemia engraftment.

Mouse study 2 (0.5x10^6^ CAR^+^ T cell dose)

Mouse study 2 was performed by the laboratory of Renata Stripecke at Medizinische Hochschule Hannover, Germany. Mouse experiments were performed in accordance with the German Animal Welfare Act and the EU-directive 2010/63 and were approved by the Lower Saxony Office for Consumer Protection and Food Safety (“Niedersächsiches Landesamt für Verbraucherschutz und Lebensmittelsicherheit, LAVES; Protocols Nr.: 33.12-42502-04-21/3791 and 33.12-42502-04-16/2347). Immunodeficient NRG mice were obtained from the Jackson Laboratory (JAX; Bar Harbor, Maine) and bred under pathogen-free conditions. 8 weeks-old mice were infused with 0.5x10^6^ Nalm-6/GFP-fLuc cells injected intravenously (i.v.) into the tail vein and control mice were injected with PBS. Four days after leukemia challenge, 0.5x10^6^ or 1x10^6^ TCR-deficient CD19-CAR T cells were infused i.v.. Those CAR T cells were generated either via targeted integration of a CAR or a *trunc*CAR into the *TRAC* or *CD3ζ* gene, respectively, or by LV gene transfer and consecutive *TRAC*-KO. All cell products were depleted of residual TCR^+^ T cells via MACS (CD3 microbeads and LD columns, Miltenyi Biotec). CAR T cells were applied freshly without prior cryopreservation on day 18 post blood collection / day 16 post non-viral gene editing. For leukemia burden assessments, mice were anesthetized using isoflurane and injected intraperitoneally (i.p.) with 2.5 mg D-Luciferin potassium salt dissolved in 100 µl PBS. 5 min after, BLI analyses were performed with the IVIS SpectrumCT apparatus (PerkinElmer, Waltham, Massachusetts) and the data was analyzed using LivingImage software (PerkinElmer). Euthanasia was performed five weeks after leukemia challenge according to the animal experiment protocols. The controls groups (tumor only, LV CAR T cells, *TRAC*-CAR T cells) of the mouse model 2 were previously published in Kath et al at Molecular Therapy Methods and Clinical Development (Figure 6) under a ‘Creative Commons Attribution - NonCommercial – NoDerivs (CC-BY-NC-ND 4.0)’ license ^2^. The published data serve as a control for this group. The *CD3ζ* -edited CAR T cells population is now published for the first time.

Mouse study 3 (0.5x10^6^ CAR^+^ T cell dose)

Mouse study 3 was performed by EPO Berlin GmbH in Berlin, Germany, in accordance with the German Animal Welfare Act and approved by local authorities (Landesamt für Gesundheit und Soziales, LaGeSo Berlin, Germany) and under the permission A0010/19 for in vivo therapy experiments. This study was similar to mouse study 1 except for the following differences. CAR T cell dose was 0.5x10^6^ TCR-deficient CD19-CAR T. CAR T cells were applied without prior cryopreservation on day 6 after blood collection / day 4 after non-viral gene transfer. The cell products were not depleted of residual TCR/CD3^+^ T cells. Mice were euthanized after individual evaluation for each mouse with BWL >20% or ethical end points reached (health score < 2).

*In vitro live cell imaging of Her2^+^ tumor cells*

HER2-positive GFP-transduced tumor cell lines were seeded one day before adding the HER2-CAR-T cells at half the density of the desired target cell number. The next day, HER2-CAR-T cells were added at a CAR^+^ Effector:Tumor ratio of 1:20. The amount of CAR^+^ T cells was adjusted to the lowest CAR rate and the cells were diluted with WT T cells to an equal cell number.

*CAST-Seq analysis of potential off-targets during CD3ζ-editing*

Genomic DNA of the samples was extracted using the NucleoSpin Tissue® kit (Macherey-Nagel). CAST-Seq analyses were conducted following the protocol previously described^3^, with some modifications: Average fragmentation size of the genomic DNA was aimed at a length of 500 bp and the generated libraries were sequenced on a NovaSeq 6000 using 2x 150 bp paired-end sequencing (GENEWIZ, Azenta Life Sciences). The bioinformatic pipeline was adjusted to improve specificity with following changes: Sites under investigation were categorized as OMT if the p-value met the cutoff of 0.005. Annotation for barcode hopping was included in the CAST-Seq algorithm, and coverage analysis has been revised to reduce the execution time by aligning the gRNAs only to the most covered regions for each site, as described for T-CAST.^4^ The CAST-Seq analysis was performed with two technical replicates from samples of two different donors. Only sites identified as significant hits in both biological replicates were considered as putative events. Results are showed in Supplementary Figure 7 and Supplementary Table S1.

**References for Supplementary Methods**

1.	Hermans IF, Silk JD, Yang J, et al. The VITAL assay: a versatile fluorometric technique for assessing CTL- and NKT-mediated cytotoxicity against multiple targets in vitro and in vivo. *J Immunol Methods*. 2004;285(1):25–40.

2.	Kath J, Du W, Pruene A, et al. Pharmacological interventions enhance virus-free generation of TRAC-replaced CAR T cells. *Mol Ther Methods Clin Dev*. 2022;25:311–330.

3.	Turchiano G, Andrieux G, Klermund J, et al. Quantitative evaluation of chromosomal rearrangements in gene-edited human stem cells by CAST-Seq. *Cell Stem Cell*. 2021;28(6):1136-1147.e5.

4.	Rhiel M, Geiger K, Andrieux G, et al. T-CAST: An optimized CAST-Seq pipeline for TALEN confirms superior safety and efficacy of obligate-heterodimeric scaffolds. *Front Genome Ed*. 2023;5:1130736.
